# Cell polarisation in a bulk-surface model can be driven by both classic and non-classic Turing instability

**DOI:** 10.1101/2020.01.29.925628

**Authors:** Johannes Borgqvist, Adam Malik, Carl Lundholm, Anders Logg, Philip Gerlee, Marija Cvijovic

## Abstract

The GTPase Cdc42 is the master regulator of eukaryotic cell polarisation. During this process the active form of Cdc42 is accumulated at a particular site on the cell membrane called the *pole*. It is believed that the accumulation of the active Cdc42 resulting in a pole is driven by a combination of activation-inactivation reactions and diffusion. It has been proposed using mathematical modelling that this is the result of diffusion-driven instability, originally proposed by Alan Turing. In this study we developed, analysed and validated a 3D bulk-surface model of the dynamics of Cdc42. We show that the model can undergo both classic and non-classic Turing instability by deriving necessary conditions for which this occurs and conclude that the non-classic case can be viewed as a limit case of the classic case of diffusion driven instability. We thoroughly investigate the parameter space. Using three-dimensional spatio-temporal simulation we predicted pole size and time to polarisation, suggesting that cell polarisation is mainly driven by the reaction strength parameter and that the size of the pole is determined by the relative diffusion.

## Introduction

*Cell division control protein 42 homolog, Cdc42* is an enzyme of the class GTPases that regulates various signalling pathways involved in cell division and cell morphology [20, 4]. It is one of the most conserved GTPases where the Cdc42 in yeast is 80% identical to that in human cells [17]. In the late G_1_-phase during the cell cycle, a sequence of events causes the accumulation of Cdc42 [5], which is the master regulator of cell division, at a specific location on the cell membrane. This location is called the *pole* which is the site where the new cell grows out and the latter process is called budding in the case of the yeast *Saccharomyces cerevisiae*. Moreover, in the cytosol Cdc42 is bound to GDP which corresponds to its inactive state while it is bound to GTP in the membrane corresponding to its active form. The conversion between these two states is catalysed by the two classes of enzymes called GEFs and GAPs. It is believed that it is the combination of these reactions of activation and inactivation along with diffusion that results in accumulation of active Cdc42.

Experimentally, the challenge with studying the activation of Cdc42 is that the concentration profile is not uniform in the cell. Thus, accounting for spatial inhomogeneities is crucial when data of the pathway is collected, however measuring two different diffusion rates simultaneously is difficult. Firstly, measuring the slow diffusion rate of active Cdc42 on the cell membrane is not trivial. Secondly, in addition to accounting for the spatial distribution of Cdc42, the activation and inactivation reaction rates should be measured as well. Usually, such rates are estimated from data of spatial averages of the concentration profiles over time which is perhaps not feasible to do in the case of the mentioned polarisation system as inhomogeneous distributions of proteins are crucial for the function of the system. On account of these experimental limitations, computational models have been developed to aid in understanding the activity of Cdc42.

Numerous mathematical models of the dynamics of Cdc42 have been developed [14, 31, 18, 35, 10, 13, 26, 7, 24, 16, 33, 9, 15, 12]. Many of these models can be reduced to a classic *activator-inhibitor* system focusing on the spatial and temporal dynamics of active and inactive Cdc42 [14, 31, 7, 26, 16, 33, 12]. An important consideration in such models is whether the accumulation of active Cdc42 in a single location is the result of a Turing-type mechanism or not. Early models of polarisation have a single spatial dimension describing either the chemical concentration along a diameter of the cell or the cell perimeter while considering the cytosol as spatially uniform. Later however, a model on the single-cell scale was developed where Turing patterns formed on the cell membrane in the presence of non-linear reactions involving another species diffusing in the cytosol [19]. It was demonstrated that with this new type of bulk-surface model, a distinct type of pattern formation mechanism was possible. In classic Turing-type systems, equal diffusion rates of the reacting species can never produce patterns, however this is no longer a necessary restriction in the bulk-surface model. It has been argued that the necessary difference in transport can be achieved by choosing the various reaction-rates to be unequal and this is referred to as non-classic Turing patterns [27, 29]. Sufficient conditions for the emergence of both classic and non-classic Turing patterns in the context of bulk-surface models have been derived and demonstrated [29, 27]. Additionally, a previous one-dimensional model of cell polarisation [24] has been extended to the bulk-surface context [8]. This model is a two-species system of active and inactive GTPases, and does not distinguish between the inactive form in the cytosol and the inactive form in the membrane. Effort has been made to extend classical 1D models into 2D [12] as well as including a number of more complicated phenomena such as a diffusion barrier on the membrane, and the presence of organelles in the cytosol.

Although bulk-surface models of polarisation is not a novel concept, most previous work has been focused on the occurrence of pattern formation and the qualitative behaviour of the models. However, little has been done in order to investigate the parameter space and the regions that give rise to classic or non-classic Turing patterns. In this work, we constructed a reaction-diffusion model of Cdc42 activation with the aim to propose an underlying mechanism of cell polarisation. We use mathematical analysis to investigate the two cases of classic and non-classic diffusion driven instability. Moreover, we conducted an analysis of the parameter space and investigated how it influences polarisation. With these results in hand we derived a necessary condition for diffusion-driven instability and proposed that the model can form patterns through both classic and non-classic Turing instability. Finally, we validated the proposed mechanisms using three-dimensional spatio-temporal simulations of the developed model. This resulted in precise conditions allowing for the formation of a single pole and determining how the parameters influence polarisation time, size of the pole and the local concentration of active Cdc42 at the pole.

## Results

### The reaction diffusion model of Cdc42 activation

To derive the reaction mechanism for the polarisation process mediated by Cdc42, we describe the cell as the interior of a three dimensional ball Ω with radius *R* where its surface Γ corresponds to the cell membrane (Fig 1B):

**Figure 1:**
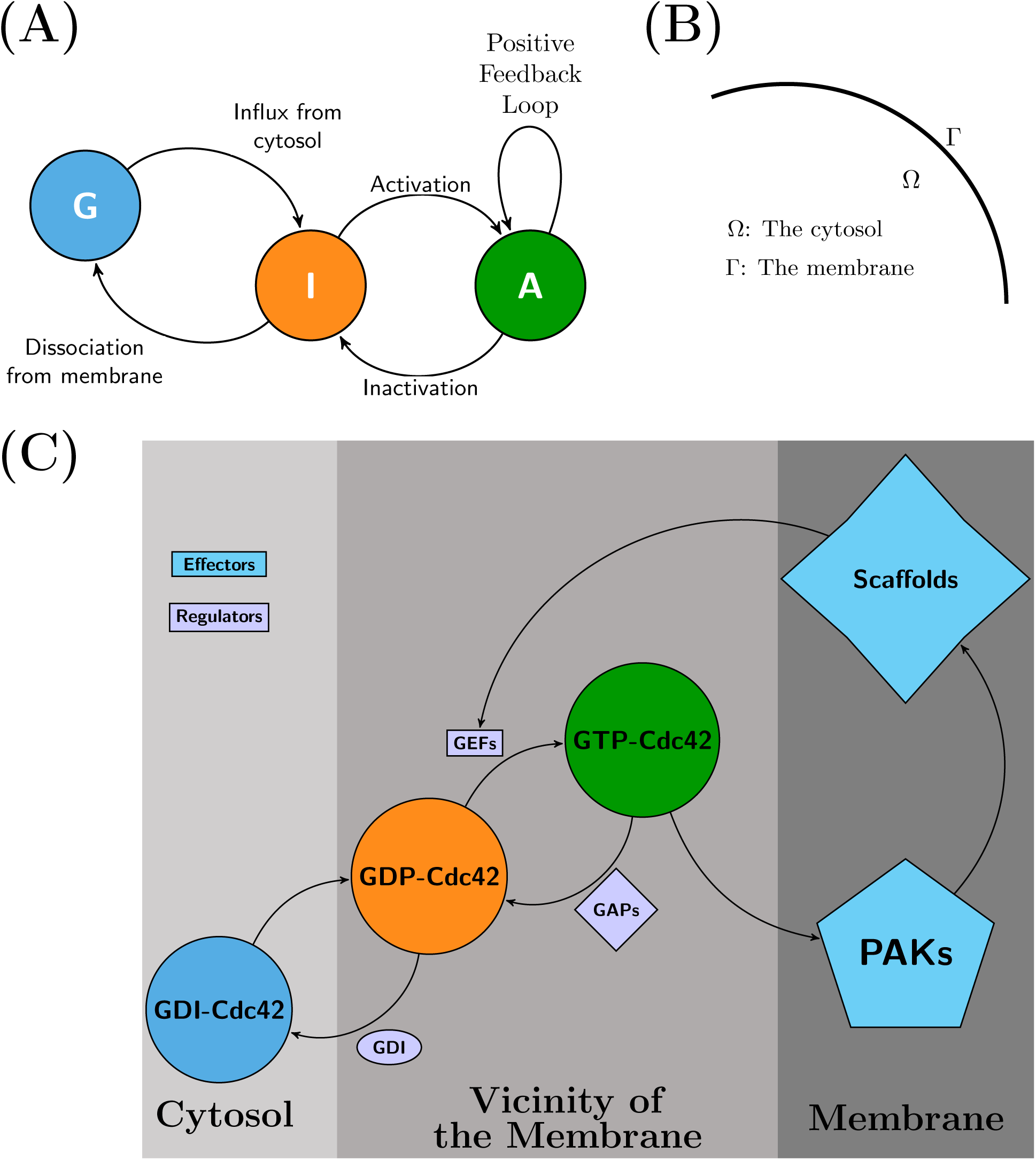
The model of Cdc42-activation. **(A)**: *The mechanism described as an expanded activator inhibitor system*. The GTPase Cdc42 is shuffled between its active state *A* (green) corresponding to the GTP-bound form and its inactive state *I* (orange) corresponding to the GDP-bound form. Also, the GDI-bound form *G* (blue) is restricted to the cytosol and is transported to the membrane which correspond to the influx of inactive form *I*. The reaction mechanism is determined by three classes of reactions: (1) Flux of inactive Cdc42 over the membrane, (2) Activation and inactivation reactions restricted to the membrane determined by GEFs and GAPs respectively and (3) The positive feedback loop mediated by PAKs and Scaffolds also restricted to the membrane. **(B)**: *Geometric domain*. By letting the membrane thickness shrink to zero, the geometric description is simplified to one domain Ω corresponding to the cytosol and one boundary Γ corresponding to the membrane. **(C)** *Biological details of Cdc42 activation*. Cdc42 has an inhibited GDI-bound form (blue), an inactive GDP-bound form (orange) and an active GTP-bound form (green). The shuffling between these forms are determined by the three regulators namely GDI, GEFs and GAPs. The active form of Cdc42, unlike the inactive, can bind to various effector molecules such as PAKs and Scaffolds which can further bind to GEFs which enhances the activation reaction through a positive feedback loop. This sub-panel is redrawn based on figures in [6].

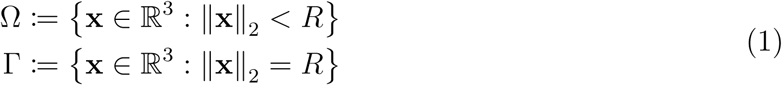

where ‖**x**‖_2_ is the Euclidian distance measure. As the cytosol Ω is comparatively large compared to the membrane it could be considered as a bulk which is subsequently linked to the two-dimensional membrane Γ. We refer to models which account for this geometric description (1) as *bulk-surface* models. In this way it is possible to expand the classic Turing framework [34] to account for reaction-diffusion models with two geometric domains which is both the most natural and more realistic than one-dimensional simplifications in the context of Cdc42.

Using this notation, we derive a *bulk-surface Activator-Inhibitor* (*AI*)-system of cell polarisation mediated by Cdc42 whose dynamics is governed by five reactions: *Influx of inactive Cdc42 from the cytosol, dissociation of inactive Cdc42 from the membrane, activation of inactive Cdc42, inactivation of active Cdc42* and *activation of Cdc42 through a positive feedback loop* (Fig 1A) (for details see Supplementary material S1.1):

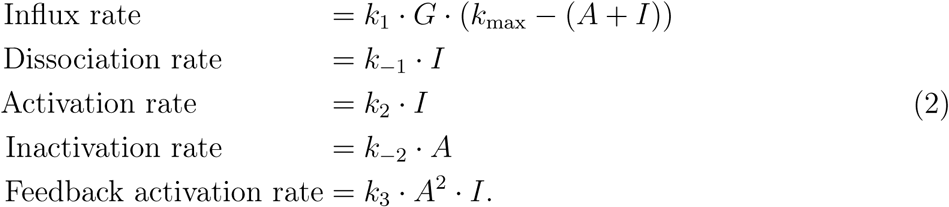

To this end, we introduce three functions

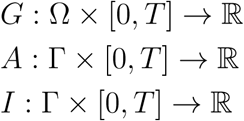

describing the concentrations of the GTP-, GDP- and GDI-bound forms respectively. Both *G* and *I* are referred to as the inactive form of Cdc42, whereas *A* corresponds to the active form.

The functions *A* and *I* are restricted to the membrane Γ while the function *G* is restricted to the cytosol Ω. The concentration in the cytosol is measured in mol*/*m^3^, and the concentrations on the membrane are measured in mol*/*m^2^.

Combining all these terms allows us to formulate a reaction diffusion model for cell polarisation mediated by Cdc42:

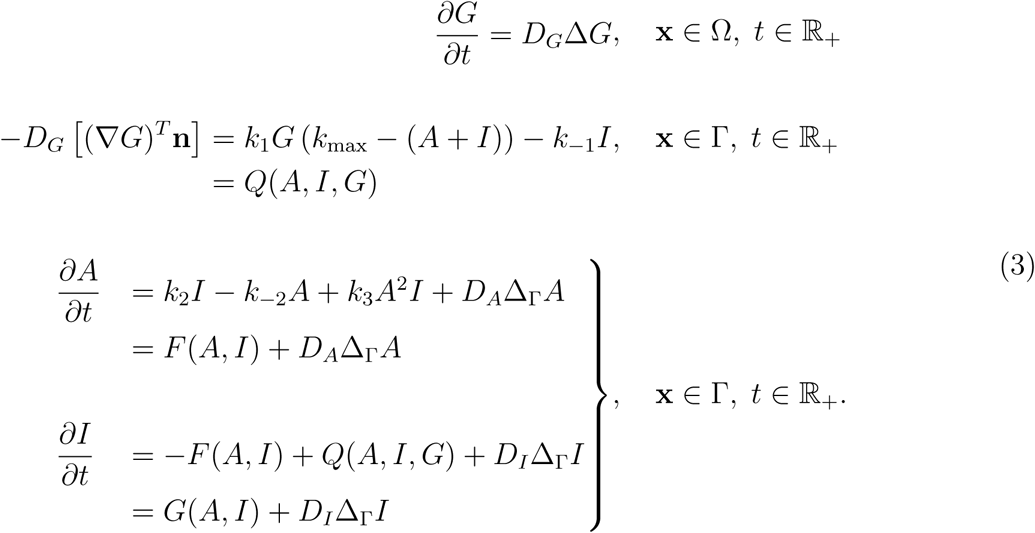

Here, the diffusion is determined by the Laplace operator 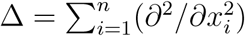 where Δ_Γ_ determines the diffusion restricted to the membrane.

The dynamics in the cytosol is entirely described by the diffusion of the GDI-bound form *G*. The flux of the inactive GDI-bound form of Cdc42 *G* from the cytosol to the membrane resulting in the influx of the membrane-bound inactive GDP-bound form of Cdc42 *I* is determined by the function *Q*. The function *Q*(*A, I, G*) is implemented as a Robin boundary condition for the GDI-bound state *G* and it is also part of the reaction term for the membrane bound GDP-bound state *I*. Moreover, the total mass of the system is conserved, independent of the choice of the functions *F* and *Q*, which can be seen by considering the temporal change of the total mass of the system, and using the partial differential equations it follow that

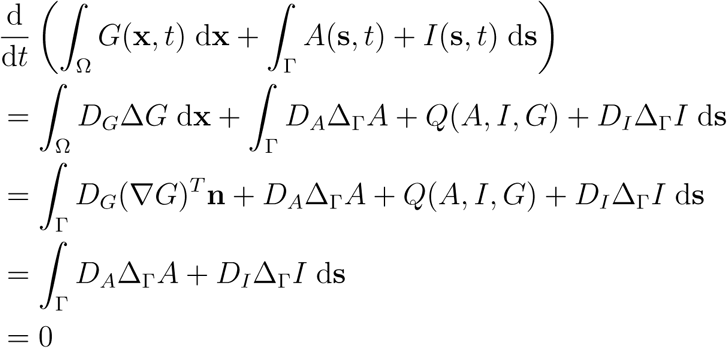

which implies that the total amount of protein is constant.

Furthermore, the model is non-dimensionalised using non-dimensional parameters similar to the ones in the classical Schnackenberg model [25] (Supplementary material S1.2), resulting in the following model structure:

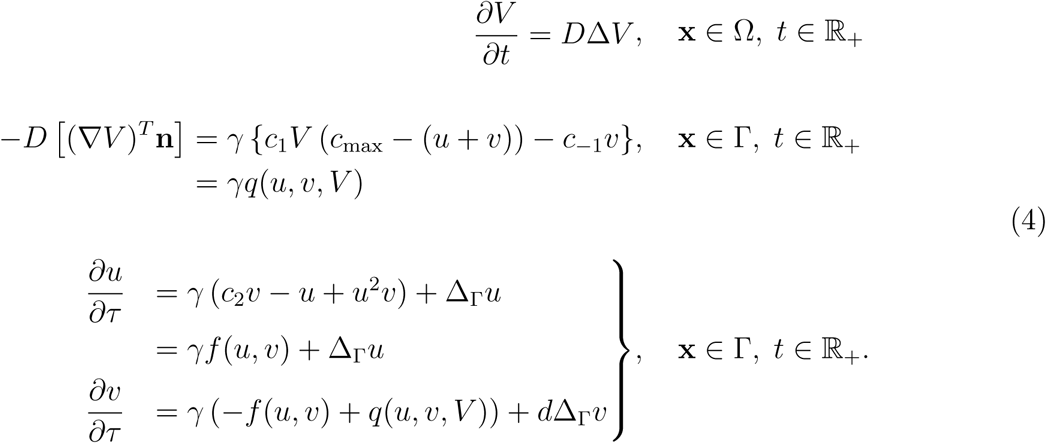

Note here that in the dimensionless model, the domain Ω corresponds to the *unit ball* and the boundary Γ corresponds to the *unit sphere*.

As a result of the non-dimensionalisation procedure, the number of parameters has been reduced from ten to eight, where the resulting dimensionless parameters are also more meaningful compared to the original ones. For example, the parameter *γ* determines the relative strength of the *reaction* part of the model compared to the *diffusion* part which implies that this parameter determines which of these forces that dominate the dynamics of the system. Moreover, all of the dynamics corresponding to the activation, inactivation and the positive feed-back loop is captured in the dimensionless parameter *c*_2_ which, in the case with dimensions, are described by the three parameters *k*_2_, *k*_−2_ and *k*_3_. The dimensionless states, variables and parameters are summarised below (Tab 1).

As the cytosolic GDI-bound state of Cdc42 diffuses much faster than the membrane bound states [6], it is possible to reduce the number of equations in (4). More precisely, the assumption that *D* → ∞implies that the concentration of the cytosolic GDI-bound state is homogeneous and in this case the mass conservation property is described by the *non-local functional V* [*u*+*v*] below

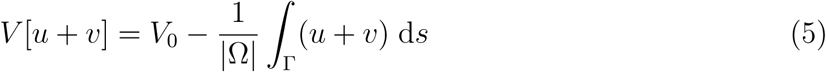

and the RD-system in (4) gets reduced to the following two-state system

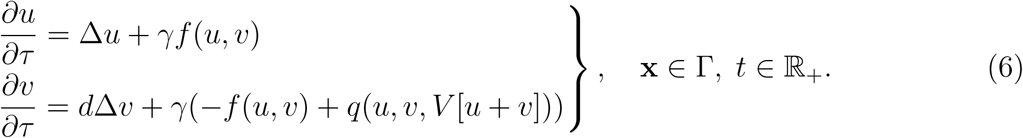

The constant *V*_0_ is the total average concentration of all three forms of Cdc42, and *V*_0_ |Ω| is the total amount of Cdc42 in the cell. The full system in (4) is solved numerically while the reduced system in (6) is analysed to determine if Turing patterns can be formed.

The model presented here builds on the framework proposed by Rätz and Röger [28, 29], where the function *f* describing the dynamics of the activation, inactivation and feedback loop of Cdc42 assumes *Michaelis–Menten* kinetics and is given by:

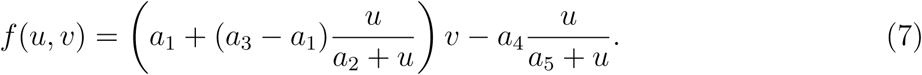

In the context of the Cdc42 model, the *Michaelis–Menten* assumption implies that the substrates would be the various states of Cdc42 while the enzymes would be the GEF’s and GAP’s.

**Table 1:** Dimensionless components of the model. The columns from left to right: the components (i.e. the states, variables and parameters), their definitions and a description of their meaning.

| Component | Definition | Description |
| --- | --- | --- |
| *The states* |  |  |
| $u$ | $u = A \cdot \sqrt{\frac{k_3}{k_{-2}}}$ | The dimensionless active form of Cdc42 (*membrane-bound*) |
| $v$ | $v = I \cdot \sqrt{\frac{k_3}{k_{-2}}}$ | The dimensionless inactive form of Cdc42 (*membrane-bound*) |
| $V$ | $V = \frac{G}{R} \cdot \sqrt{\frac{k_3}{k_{-2}}}$ | The dimensionless GDI-bound Cdc42 (*cytosolic*) |
| *The variables* |  |  |
| $\tau$ | $\tau = \frac{D_A t}{R^2}$ | The dimensionless time (S14) |
| $\mathbf{x} \in \Omega_R$ | $\mathbf{x} \leftarrow \frac{1}{R} \mathbf{x}$ | The dimensionless spatial variable (S14) |
| *The parameters* |  |  |
| $\gamma$ | $\gamma = \frac{R^2 k_{-2}}{D_A}$ | The parameter has three interpretations [25, page 78]: **(1.)** $\gamma$ is proportional to the area of the domain $\Omega$ in (1). **(2.)** $\gamma$ is the relative strength of the two reaction terms determined by $f(u, v)$ and $g(u, v)$ in (4). **(3.)** An increase in $\gamma$ corresponds to an equivalent decrease in the diffusion coefficient $d$. |
| $c_1$ | $c_1 = \frac{k_1}{k_{-2}} \sqrt{\frac{k_{-2}}{k_3}}$ | The relative influx of inactive Cdc42 from the cytosol |
| $c_{\max}$ | $c_{\max} = k_{\max} \sqrt{\frac{k_3}{k_{-2}}}$ | The maximum amount of membrane bound Cdc42, i.e. $(u + v) \leq c_{\max}$ |
| $c_{-1}$ | $c_{-1} = \frac{k_{-1}}{k_{-2}}$ | The relative dissociation of inactive Cdc42 from the membrane |
| $c_2$ | $c_2 = \frac{k_2}{k_{-2}}$ | The relative activation rate of Cdc42 |
| $d$ | $d = \frac{D_I}{D_A}$ | The relative diffusion rate of inactive Cdc42 (*diffusion in the membrane* $\Gamma$) |
| $D$ | $D = \frac{D_G}{D_A}$ | The relative diffusion rate of GDI-bound Cdc42 (*diffusion in the cytosol* $\Omega$) |
| $V_0$ | $V_0 = G_0 \cdot \sqrt{\frac{k_3}{k_{-2}}}$ | The initial amount of dimensionless GDI-bound Cdc42 |

However, as Cdc42 itself is an enzyme it is more reasonable to assume that its intracellular concentration is in the same order of magnitude as that of the GAP’s and the GEF’s. Therefore, we describe the dynamics of Cdc42 governed by the activation, inactivation and the feedback loop with the simpler structure

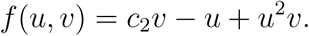

Here, the parameter *c*_2_ describing the relative activation rate of Cdc42 is well-motivated by the litterature (Fig 1). In contrast to the previous framework, involving a larger number of arbitrary parameters [28, 29], the simplicity of our reaction term ensures that our approach can qualitatively model cell polarisation as each parameter has a concrete meaning. For example, a large value of the parameter *c*_2_ relative to the value of *c*_−1_ implies a high activity of the activation-inactivation module monitored by GEF’s, GAP’s and the positive feedback-loop relative to the dissociation from the membrane implying the attachment of GDIs to the inactive form of Cdc42.

### The bulk-surface model can form patterns through both classic and non-classic Turing-patterns

Given the derived model, we can answer three fundamental questions. Does a unique solution to the RD system in (6) determined by the initial conditions *exist* ? If so, are the solutions physically realistic, meaning that they give rise to non-negative and bounded concentrations of *u* and *v*? If this is the case, can the model undergo diffusion driven instability and thereby form patterns? To answer these questions, we define the *homogeneous* system of the reduced model in (6) as follows

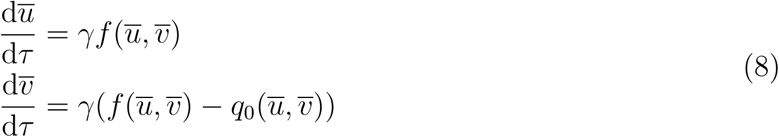

where the states 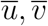 can be viewed as spatial averages. Here, the function 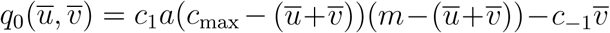 where *m* = *V*_0_*/a* is obtained as a consequence of the spatial averaging as the functional *V* in (5) is replaced by 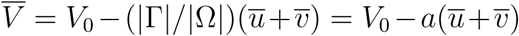 where *a* = 3 is the ratio between the area of the unit sphere divided by the volume of the unit ball. Moreover, as we are interested in non-negative states, we require that 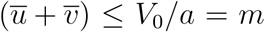 in (8) which implies 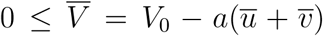 where the expression for the cytosolic component 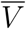 stems from the mass conservation functional. In addition, the total amount of membrane-bound proteins is also constrained by *c*_max_, i.e. 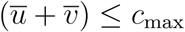. Therefore, physically reasonable states corresponding to 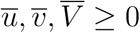 lie in the region *A* in the (*u, v*)-state space defined as follows

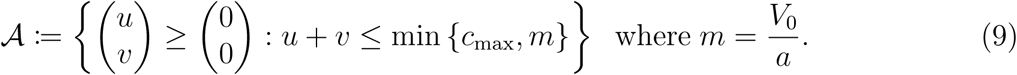

#### Existence of a unique solution

The existence of solutions to RD models is not guaranteed for all continuous reaction functions *f* and *q*. To this end, we prove the existence of a unique solution (Thm 1) to the reduced model in (6) if we choose the initial conditions from the previously defined region *A* in (9) (Supplementary Material S1.3).

##### Theorem 1

(**Existence of a unique solution on the unit sphere in global time**). *Consider the RD model in* (6) *with initial conditions u*(**x**, 0) = *u*_0_(**x**) *and v*(**x**, 0) = *v*_0_(**x**) *chosen such that* (*u*_0_(**x**), *v*_0_(**x**)) ∈ *A* ∀ **x** ∈ Γ *where the region A is defined in* (9). *Then, there exists a unique solution to* (6) *global in time.*

#### Physical validity of the model

We have already established one physical property, namely *mass conservation*, which governs the system. However, this property follows from the *structure* of the model implying that *any* choice of the reaction terms *f* and *q* results in a model with this property. Also, the fact that the mass of the states *u, v* and *V* is conserved does not prevent non-physical behaviour of the solutions such as negative concentrations of the involved proteins,e.g. *u*(**x**, *τ*) ≤ 0 for some coordinate **x** ∈ Γ at some time *τ*. To this end, we prove (Thm 2) that the region *A* in (9) is a *trapping region* meaning that if the initial conditions are chosen within this region then we will always have non-negative solutions at all times (Supplementary Material S1.4). Interestingly enough, from this follows another theoretical result (Corollary 1) which states that the domain Γ can deviate from the unit sphere and we will still have a unique solution to the RD model in (6) global time (for details see Supplementary Material S1.4). The latter result implies that with our model, i.e. choice of *f* in combination with *q*, the shape of the membrane can deviate from the sphere and the model will still have a solution that is uniquely determined by its initial conditions.

##### Theorem 2

(**Existence of a trapping region**). *The region A* ⊂ ℝ^2^ *in* (9) *is a trapping region meaning that the trajectories of the solutions to the homogeneous system in* (8) *can never leave the region for initial conditions* (*u*(0), *v*(0)) = (*u*_0_, *v*_0_) ∈ *A*.

##### Corollary 1

(**Existence of a unique solution on any open interval in** ℝ^*n*^ **in global time**). Consider the RD model in (6) where the domain Γ ⊂ ℝ^*n*^ for any *n* ∈ ℕ_+_ is changed to an open, bounded and regular domain with Neumann (i.e. zero-flux) boundary conditions. If the initial conditions *u*(**x**, 0) = *u*_0_(**x**) and *v*(**x**, 0) = *v*_0_(**x**) are chosen such that (*u*_0_(**x**), *v*_0_(**x**)) ∈ *A* ∀**x** ∈ Γ where the region *A* is defined in (9), then this problem has a unique solution global in time.

#### Diffusion driven instability

The underlying idea behind pattern formation caused by diffusion-driven instability pattern formation was originally formulated by Turing [34], and it entails a switch in stability in the sense of linear stability analysis. The phenomena depends on the reaction terms, e.g. *f* and *q*, having the capacity to allow for the existence of a stable steady state (*u*^∗^, *v*^∗^) to the homogeneous ODE-system obtained by neglecting diffusion. When diffusion is introduced, *classic* diffusion-driven instability occurs when a *stable node* in the homogeneous system is switched to an *unstable node* in the inhomogeneous system. In the *non-classic* diffusion-driven instability [28, 29] this takes place when a steady-state (*u*^∗^, *v*^∗^) transitions from a *stable node* in the homogeneous system to a *saddle point* in the inhomogeneous system. The exact formulation of the mathematical conditions for these two cases are presented in the Supplementary material (section S1.5). It is important to emphasise that both cases rely on the existence of a steady-state with the capacity of switching stability, and it is thus crucial to find rate parameters ensuring this fundamental requirement. To this end, we have mathematically proven that the model has steady states, and we have derived a necessary condition ensuring the stability of a specific steady state. (Thm 3) (Supplementary material S1.6).

##### Theorem 3

(**Existence and characterisation of steady states**). *The system in* (8) *has either 2, 4 or 6 positive steady-states within the first quadrant of the* (*u, v*)*-state space. The following condition is necessary*

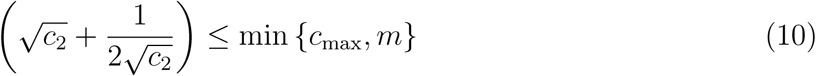

*to ensure the existence of a steady-state* (*u*^∗^, *v*^∗^) ∈ *A in the trapping region in* (9). *Lastly, a steady-state* (*u*^∗^, *v*^⋆;^) ∈ *A allowing for diffusion-driven instability satisfies the following bounds* 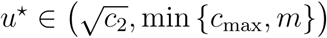.

The necessary condition in (10) implies that the activation-inactivation parameter *c*_2_ is constrained by the maximum concentration of membrane bound species, *c*_max_, and the average total concentration, *V*_0_. Furthermore, we show that there is always one such steady-state (possibly more) which could potentially give rise to patterns and that it always lies in the interval 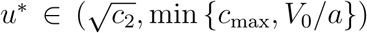. These bounds indicate that using the rate parameter *c*_2_, cor-responding to the activation-inactivation reactions, it is possible to formulate a lower bound on the steady-state. Combining the maximum concentration of membrane-bound species *c*_max_ with the total average concentration *V*_0_ of the cytosolic state the upper bound can be formulate. Note that these conditions merely ensure the existence of steady-states, and to ensure the emergence of patterns the exact steady-state satisfying the mathematical conditions in the stability analysis must be found.

#### Numerical mappings of the parameter space indicate that the nonclassic case of diffusion driven instability is a limiting class of the classic case

Next, we investigate the steady-states by numerical exploration of the parameter space to determine whether they satisfy the classic or non-classic Turing conditions derived by Röger and Rätz [28, 29]. To this end, we investigated a large set of kinetic parameters by mapping out the (*c*_−1_, *c*_2_) and (*c*_1_, *c*_2_) cross sections of the parameter space. For each point in these cross sections, two kinetic parameters are varied while the remaining parameters are kept fixed. We observed that for a large relative diffusion, *d*, the set of parameters enabling pattern formation is larger in the classic than the non-classic case (Fig 2). It is worth emphasising that the region where the Turing conditions are met for *d* = 5, is a subset of the region for any larger *d* and in the figures, we have chosen to plot the regions in layers, with the lowest *d* shown on top. Also, the non-classic region is special in the sense that the classical Turing conditions can never be met when *d* = 1. This implies that the set of parameters that allow for symmetry breaking increases with the relative diffusion, *d*, in both cases (Fig 2). Conversely, when the the relative diffusion decreases the region of the parameter space allowing for diffusion-driven instability decreases as well, and in fact the non-classic case can be viewed as the classic case in the limit *d* → 1. Note here, that the non-classic case is independent of the relative diffusion *d* and in the case where *d* = 1 the system can only form patterns through non-classic diffusion driven instability. However, as soon as *d >* 1 symmetry breaking can occur through both mechanisms and when the relative diffusion is large the classic region of the parameter space in larger than the non-classic counterpart.

**Figure 2:**
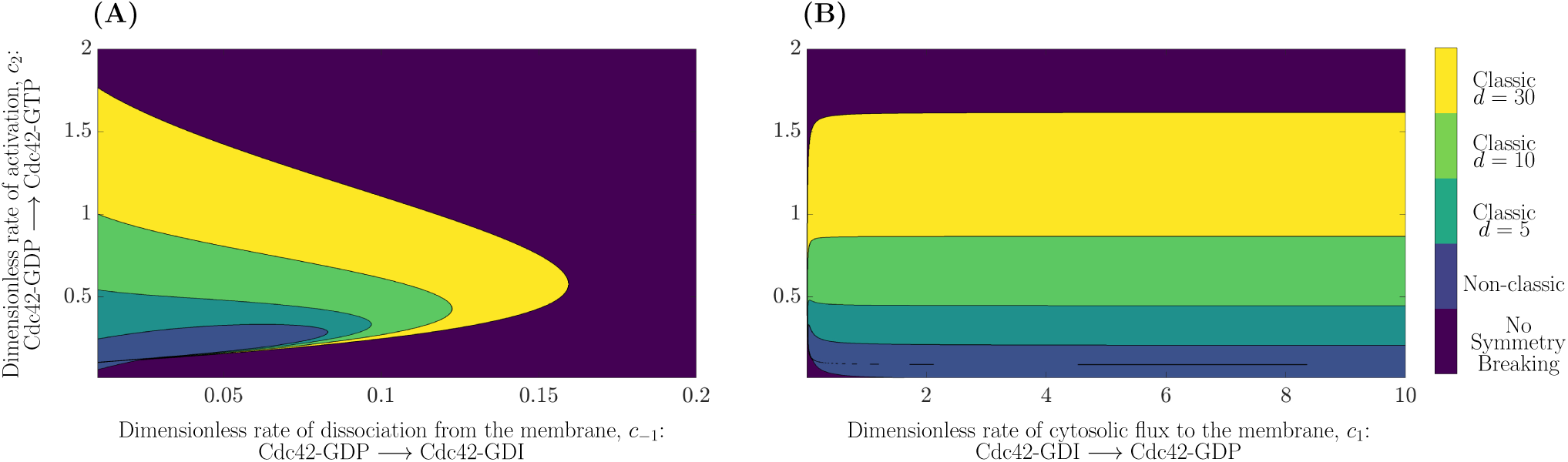
The parameter space for classic and non-classic diffusion-driven instability. The parameter space is divided into five regions indicated by the colour bar: *Classic Turing instability* with *d* = 30 (yellow), *Classic Turing instability* with *d* = 10 (light green), *Classic Turing instability* with *d* = 5 (green blue), *Non-classic Turing instability* (light blue) and *No Symmetry Breaking* (dark blue). **(A)** the (*c*_−1_, *c*_2_)− plane with a fixed value of *c*_1_ = 0.05. **(B)** the (*c*_1_, *c*_2_)−plane with a fixed value of *c*_−1_ = 0.05. The overall parameters in both cases are *V*_0_ = 6.0 and *c*_max_ = 3.0 (note that the non-classic case is independent of *d*).

Further, we observe that the relative activation rate *c*_2_ must be larger than the relative dissociation rate *c*_−1_ in order to allow classic diffusion driven instability (Fig 2A). For all tested values of the relative diffusion, the relative activation rate *c*_2_ is approximately five to ten times larger than *c*_−1_. Additionally, within a range of relative activation rates *c*_2_ the phenomena of diffusion driven instability is *independent* of the relative influx rate *c*_1_ (Fig 2B). This conclusion holds true for larger values of the relative cytosolic flux than *c*_1_ = 10 although the parameter space is only illustrated up to this value. Note that a general result from both parameter planes is that classic Turing instability occurs for higher relative activation rates *c*_2_ compared to the non-classic case.

Provided these theoretical results, we next modelled cell polarisation using numerical simulations. We are interested in a specific pattern namely the formation of a single pole corresponding to a single circular spot of active Cdc42 on the cell membrane.

#### Cell polarisation can be modelled through both classic and non-classic Turing-patterns

Cell polarisation can be modelled by both cases of diffusion driven instability (Fig 3). Although the time evolution of the concentration profiles are slightly different, the final patterns are qualitatively very similar for the two cases. The classic case (Fig 3A) forms a circular pole directly while the non-classic case (Fig 3B) initially forms elongated pattern which gradually transitions into a pole. Given that the model can simulate cell polarisation, we further investigate the impact of the kinetic rate constants *c*_1_, *c*_−1_ and *c*_2_, the relative diffusion *d* and the reaction strength *γ* on cell polarisation. To this end, we define three quantitative measures of polarisation: the size of the pole, the time to polarisation as well as the maximum and minimum concentration of active Cdc42 in order to quantify the effect of altering the various parameters. In the interest of comparing the previously mentioned measures between different cases of diffusion driven instability as well as for different sets of parameters, we have implemented a “pole recognition”-algorithm (Supplementary Material S3) which terminates the simulation when a pole has been formed.

**Figure 3:**
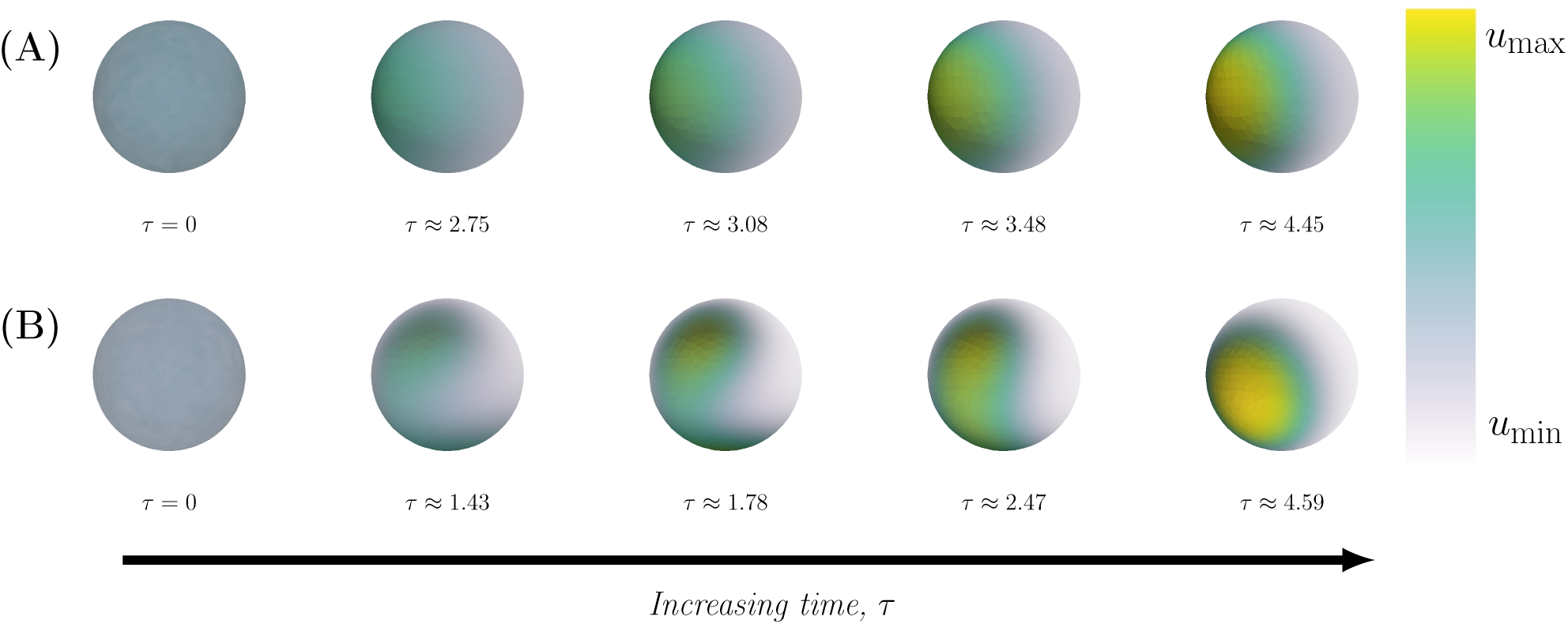
The time evolution of a pattern. The time-evolution of the concentration profiles for two sets of parameters corresponding to classic and non-classic respectively. **(A)** *Classic*: The parameters are (*c*_−1_, *c*_2_) = (0.02, 0.45) where the time points, from left to right, are *τ* = 0, *τ* ≈ 2.75, *τ* ≈ 3.08, *τ* ≈ 3.48 and *τ* ≈ 4.45. **(B)** *Non-classic*: The parameters are (*c*_−1_, *c*_2_) = (0.01, 0.20) where the time points, from left to right, are *τ* = 0, *τ* ≈ 1.43, *τ* ≈ 1.78, *τ* ≈ 2.47 and *τ* ≈ 4.59. In both cases, the overall parameters are: *c*_1_ = 0.05, *V*_0_ = 6.0, *c*_max_ = 3.0, *a* = 3, *d* = 10 and *γ* = 25.

#### The effect of the relative influx, the disassociation and activation rates of Cdc42 on cell polarisation

The final patterns for different kinetic parameters are qualitatively similar but quantitatively different (Supplementary Material Fig S4). Both the classic (Supplementary Material Fig S4A) and the non-classic (Supplementary Material Fig S4B) case form a single pole for different parameters within the (*c*_−1_, *c*_2_) − space (Fig 2A). However, from a quantitative perspective the time it takes to form a pole, *τ*_final_, differs for different sets of kinetic parameters. For instance, in the classic case it varies from *τ*_final_ ≈ 6.5 to *τ*_final_ ≈ 19.1 (Supplementary Material Fig S4A), while in the non-classic case it varies from *τ*_final_ ≈ 2.88 to *τ*_final_ ≈ 9.0 (Supplementary Material Fig S4B). Similarly, the maximum concentration of active Cdc42 *u*_max_ is different for different kinetic parameters. In the classic case, it varies from *u*_max_ = 1.54 to *u*_max_ = 3.65 (Supplementary Material Fig S4A) while in the non-classic case it varies from *u*_max_ = 3.86 to *u*_max_ = 4.02 (Supplementary Material Fig S4B). Similar conclusions can be drawn in the case of different parameters in the (*c*_1_, *c*_2_)-plane (Supplementary Material Fig S5).

#### The effect of increasing the relative diffusion on cell polarisation

A single pole is formed in both the classic and non-classic case (Fig 4) for all investigated cases of increasing relative diffusion *d*. An increase of the relative diffusion causes a decrease of the size of the pole, a decrease of the time to polarisation and an increase of the maximum (local) concentration of active Cdc42 in the pole (Fig 5 and Supplementary Material Fig S6). We did not observe any significant difference, both qualitatively (Fig 4) and quantitatively (Fig 5), between the two cases of diffusion driven instability.

**Figure 4:**
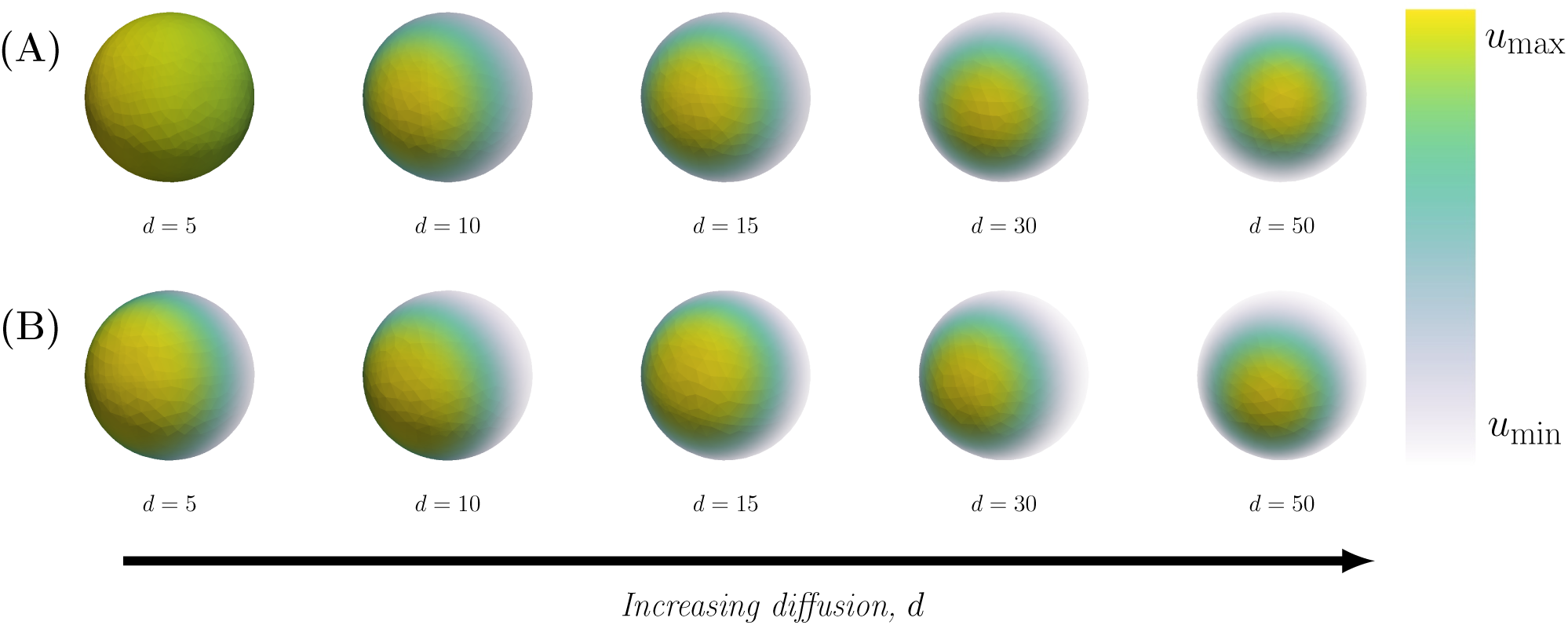
Final patterns for increasing relative diffusion with a relative scale. The final patterns for increasing relative diffusion *d* are displayed in two cases, namely classic and non-classic diffusion driven instability. In both cases, the final time when the pattern is formed *τ*_final_ and the maximum and minimum concentrations of active Cdc42 *u*_max_ and *u*_min_ are calculated as functions of the kinetic rate parameters. **(A)** *Classic*: The overall parameters are (*c*_1_, *c*_−1_, *c*_2_) = (0.05, 0.04, 0.45) with specific parameters (from left to right): no pattern is formed for (*d, τ*_final_, *u*_max_, *u*_min_) = (5, 15, 20.89, 1.34, 1.11), (*d, τ*_final_, *u*_max_, *u*_min_) = (10, 4.44, 3.65, 0.40), (*d, τ*_final_, *u*_max_, *u*_min_) = (15, 3.65, 4.85, 0.30), (*d, τ*_final_, *u*_max_, *u*_min_) = (30, 2.65, 7.42, 0.20) and (*d, τ*_final_, *u*_max_, *u*_min_) = (50, 2.17, 9.83, 0.15). **(B)** *Non-classic*: The overall parameters are (*c*_1_, *c*_−1_, *c*_2_) = (0.05, 0.03, 0.15) with specific parameters (from left to right): (*d, τ*_final_, *u*_max_, *u*_min_) = (5, 4.0, 2.77, 0.17), (*d, τ*_final_, *u*_max_, *u*_min_) = (10, 4.38, 4.18, 0.11), (*d, τ*_final_, *u*_max_, *u*_min_) = (15, 2.87, 5.29, 0.09), (*d, τ*_final_, *u*_max_, *u*_min_) = (30, 1.95, 7.77, 0.06) and (*d, τ*_final_, *u*_max_, *u*_min_) = (50, 1.96, 10.166, 0.04). In both cases, the overall parameters are: *V*_0_ = 6.0, *c*_max_ = 3.0, *a* = 3 and *γ* = 25.

**Figure 5:**
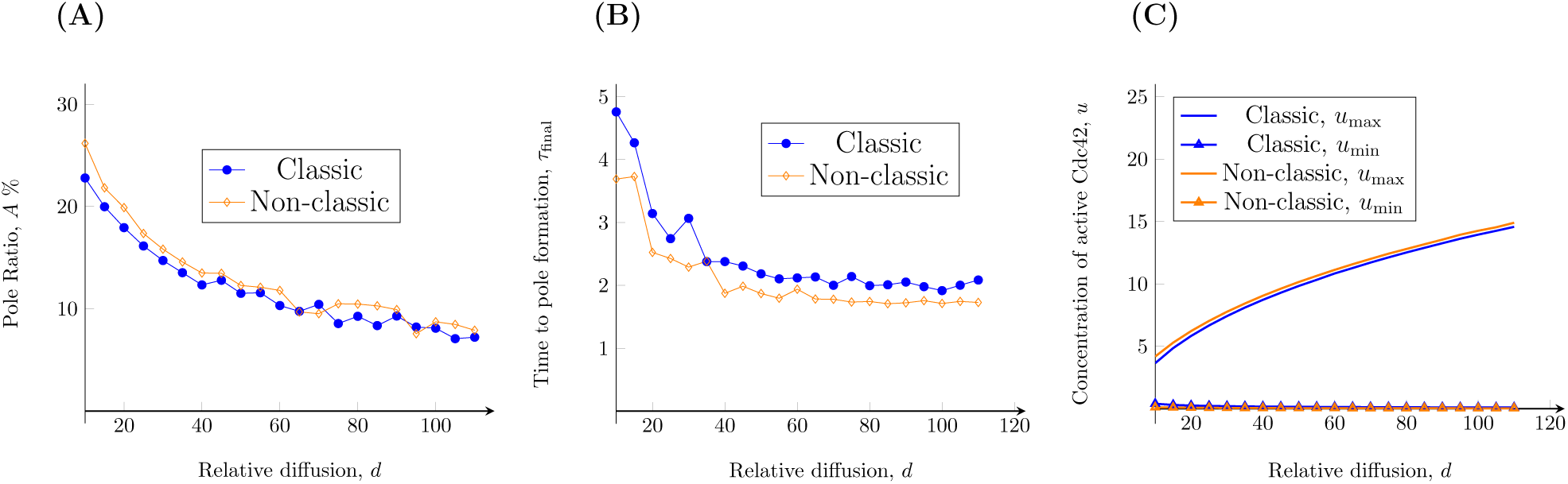
Quantitative measures as functions of an increasing relative diffusion *d*. The figure illustrates how the relative diffusion *d* influences (**A**) the size of the pole, (**B**) the time to polarisation and (**C**) the maximal and minimal values of *u* on the cell membrane.

#### The effect of increasing the relative reaction strength on cell polarisation

The number of poles increases with an increasing relative reaction strength *γ* in both the classic and non-classic case (Fig 6). The number of poles has been calculated for a large range of the parameter *γ* (Fig 7D) ranging from one single pole up to five poles. We believe that the random noise in the initial conditions are the reason for the fluctuations around the transitions between the number of poles. In addition, the relative reaction strength *γ* has no effect on the relative pole-size (Fig 7A) which varies around 25%. Again, these variations are almost certainly due to the random fluctuations in the initial data. Also, we observe that there is no clear relationship between *γ* and other quantitative measures such as the area of the pole relative to the surface area of the membrane (Fig 7A), or the maximum and minimum concentration of active Cdc42 *u* (Fig 7C). Similarly, this holds true for the classic case with respect to the time to polarisation (Fig 7B) while this property seems to increase slightly with the relative reaction strength *γ* in the non-classic case. Thus, the two cases yield different predictions when it comes to the time to polarisation.

**Figure 6:**
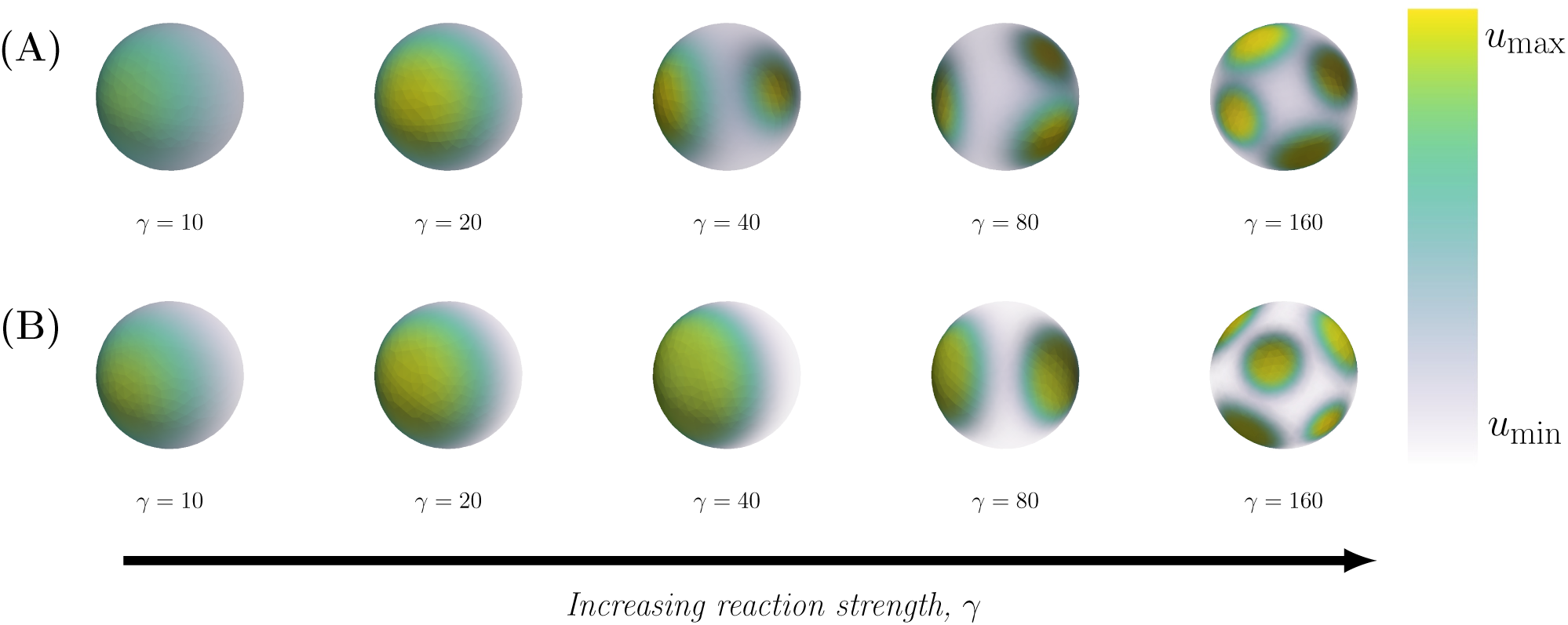
Final patterns for increasing relative reaction strength. The final patterns for increasing relative reaction strength *γ* are displayed in two cases, namely classic and non-classic diffusion driven instability. In both cases, the maximum and minimum concentrations of active Cdc42 *u*_max_ and *u*_min_ are calculated as functions of the kinetic rate parameters. **(A)** *Classic*: The overall parameters are (*c*_1_, *c*_−1_, *c*_2_) = (0.05, 0.04, 0.45) with specific parameters (from left to right): (*γ, τ*_final_, *u*_max_, *u*_min_) = (10, 10.81, 3.02, 0.49), (*γ, τ*_final_, *u*_max_, *u*_min_) = (20, 5.06, 3.62, 0.40), (*γ, τ*_final_, *u*_max_, *u*_min_) = (40, 5.86, 3.75, 0.44), (*γ, τ*_final_, *u*_max_, *u*_min_) = (80, 4.63, 3.80, 0.40) and (*γ, τ*_final_, *u*_max_, *u*_min_) = (160, 8.92, 3.74, 0.40). **(B)** *Non-classic*: The overall parameters are (*c*_1_, *c*_−1_, *c*_2_) = (0.05, 0.03, 0.15) with specific parameters (from left to right): (*γ, τ*_final_, *u*_max_, *u*_min_) = (10, 4.66, 3.96, 0.15), (*γ, τ*_final_, *u*_max_, *u*_min_) = (20, 2.77, 4.21, 0.11), (*γ, τ*_final_, *u*_max_, *u*_min_) = (40, 4.41, 4.00, 0.11), (*γ, τ*_final_, *u*_max_, *u*_min_) = (80, 11.28, 4.15, 0.12) and (*γ, τ*_final_, *u*_max_, *u*_min_) = (160, 44, 4.48, 0.11). In both cases, the overall parameters are: *V*_0_ = 6.0, *c*_max_ = 3.0, *a* = 3 and *d* = 10.

**Figure 7:**
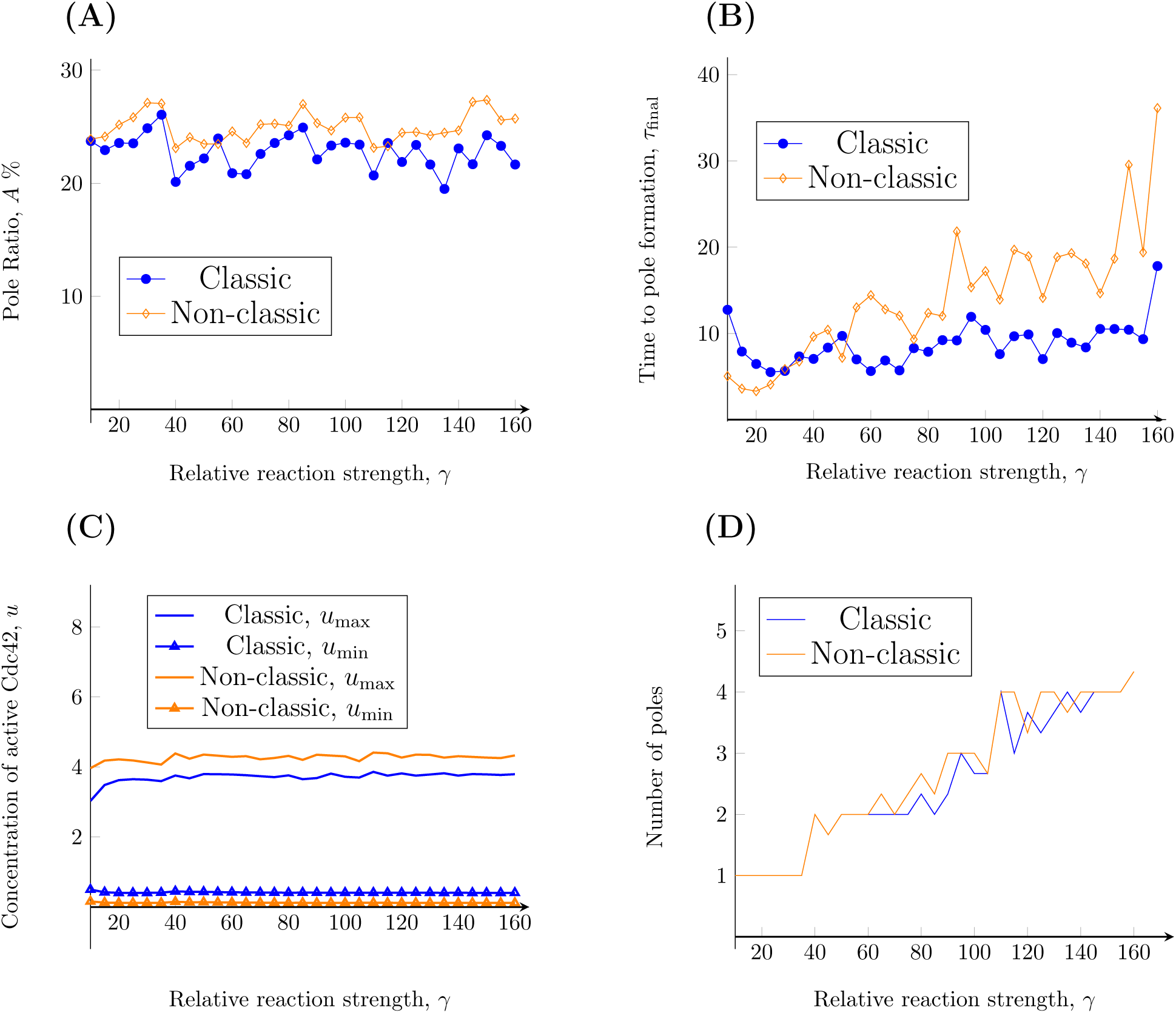
Quantitative measures as functions of increasing *γ*. The figure illustrates how the relative reaction strength *γ* influences (**A**) the size of the pole, (**B**) the time to polarisation, (**C**) the maximal and minimal values of *u* on the cell membrane, as well as (**D**) the number of poles.

## Discussion

Cell polarisation is one of the most well studied symmetry-breaking events in biology using both experimental and theoretical approaches. Yet, it still remains largely unknown how complex, intertwined, and highly dynamic protein interactions control cell polarity. In this study, we constructed, analysed and verified a bulk-surface model of Cdc42-mediated cell polarisation. The analysis of the model resulted in a mathematical theorem showing the existence of multiple steady-states. In addition, a necessary condition for diffusion-driven instability was derived. Using a thorough numerical investigation of the parameter space, we have shown that the model can form patterns by means of both classic and non-classic Turing instability. Also, the simulations highlighted the connection between these two mechanisms where the non-classic case can be viewed as the classic case for equal diffusion rates of the membrane bound species. Lastly, we validated the theoretical results by showing that both these mechanisms can sustain pattern formation. Using simulations, we propose that cell polarisation is mainly driven by a low value of the reaction strength parameter *γ*, that the size of the pole is determined by the relative diffusion *d* and that the effect of changing the kinetic parameters is quantitative rather than qualitative.

Within our bulk-surface formulation of the model, the membrane bound reaction terms and the non-dimensionalisation are novel. The choice of the geometrical description that includes both the membrane and the cytosol in combination with adding the cytosolic GDI-bound form of Cdc42 to the model [28, 29] increases the level of realism. Previous models of the “wave-pinning” type [15, 8, 24, 16] have only focused on the two membrane-bound species and assumed mass-conservation in the membrane, with one exception ([8]) that includes a cytosolic state but no extra reactions associated with it. We argue that from a biological perspective this is not entirely plausible as there is a fast-moving cytosolic state of Cdc42 that contributes to the transfer and dissociation reactions at the membrane. Furthermore, the introduced minimal reaction term *f* governing the activation-inactivation reactions is biologically motivated, where each term has a concrete meaning in terms of reaction rates. The non-dimensionalisation procedure implemented in the course of this work resulted in the derivation of biologically meaningful parameters *γ* corresponding to relative reaction strength and the activation parameter *c*_2_ corresponding to all membrane bound reactions governing activation, inactivation and the feed-back loop.

Our exhaustive analysis of the parameter space shows that the relative activation rate *c*_2_ is higher than the relative dissociation rate from the membrane *c*_−1_ in the classic compared to the non-classic case where these two rates are more similar (Fig 2). Biologically, the relative size between these parameters can be viewed as a kinetic “tug-of-war” between the two stable states of Cdc42, namely the cytosolic GDI-bound form and the membrane-bound GTP-bound form (Fig 1C). Also, the investigation of the parameter space reveals that for certain activation rates *c*_2_, any value of the cytosolic flux to the membrane *c*_1_ allows for diffusion driven instability in both cases (Fig 2B). This suggests that a pattern will form independent of the cytosolic flux of GDI-bound Cdc42 to the membrane. Furthermore, a general conclusion drawn by studying the Turing parameter space is that in both the classic and non-classic case the activation rate *c*_2_ is larger in the former compared to the latter. In addition, our results suggest that the non-classic diffusion driven instability is a special case of the classic one in the limit when *d* → 1 (Fig 2).

Perhaps the most interesting result of this work is that cell polarisation can be modelled by both classic and the non-classic Turing-patterns. This was demonstrated using numerical simulations, where we first showed that patterns can be formed through both mechanisms (Fig 3). However, the time-scales and dynamics of the two cases differ indicating that the mechanisms are different. The sensitivity of the final pattern in the two cases (supplementary Material Fig S4) with respect to variations in the kinetic parameters showed that the effect is quantitative rather than qualitative. More precisely, a mere change of parameters in the (*c*_−1_, *c*_2_)-plane does not alter the qualitative behaviour as a single pole is formed, however quantitative measures such as the time it takes to form the pole *τ*_final_ or the maximum concentration of active Cdc42 *u*_max_ are different for different kinetic parameters. In a similar investigation of the parameter space [32], the existence of Turing patterns was investigated for 2-species and 3-species systems with a Hill function governing the interaction between the species. It was concluded that a large number of interaction topologies are capable of producing Turing patterns, but that they were not robust to parameter changes. Our results show that our model could be considered robust, since the parameter regions for which it produces Turing patterns is large. Nevertheless, the comparison between the two cases is not straightforward since the notion of robustness greatly depends on the parameter ranges in which the stability in investigated. This is also affected by, for example, the implementation of non-dimensionalisation which is not done in [32].

In addition, we showed that the size of the pole, the time to polarisation and the maximum concentration *u*_max_ are influenced by the relative diffusion *d* (Fig 4). This presents an opportunity for new experimental studies and for connecting the simulations of the bulk-surface models to data as a measure, however crude, of the size of the pole (for example as a percentage of the entire surface of the cell) that can be used to estimate the relative diffusion. This methodology for estimating the relative diffusion is consequential as it is currently not possible to differentiate between the three states of Cdc42 by using fluorescent markers and it is thereby not possible to estimate the relative diffusion *d* of the two membrane bound species. Lastly, we showed that the key parameter determining the number of poles is the strength of the reaction term *γ* (Fig 6). More precisely, one pole is formed for values of *γ <* 40 while numerous poles are formed for larger values, suggesting that the two classical parameters in reaction-diffusion models, *γ* and *d*, are consequential in the context of cell polarisation. These parameters govern the number of allowed wave numbers [25], where the smaller the value of the parameters *γ* the fewer wave numbers contribute to the pattern formation and vice versa. This is in agreement with our simulations showing that the number of poles increases with *γ* (Fig 7). In our model, this parameter is proportional to the the surface of the cell (Tab 1). Thus, as the size of the cell increases so does the number of poles which is in agreement with previous studies [30]. This indicates that cell polarisation, i.e. the formation of a single spot corresponding to a pole, can be achieved for both mechanisms as the formation of this particular pattern is dependent of the relative strengths of the reaction part *γ* and the diffusion part *d*. Thus, it is not qualitatively possible to rule out either the classic or the non-classic cases based on the patterns formed as both mechanisms can form a pole for low values of *γ*. However, it might be possible to quantitatively distinguish between the cases by studying the concentration profiles over time and comparing the time it takes for the patterns to be formed. Nevertheless, this poses experimental challenges as it is hard to connect high qualitative three-dimensional data based on microscopy with numerous images over time.

Understanding the underlying mechanisms of cell polarisation can shed light on many fundamental processes governing cell division and cell differentiation both during normal development and in the context of disease. Building, analysing and simulating spatio-temporal models like the one proposed in this work can provide insight into mechanistic details as well as guide further experimental design. In the context of the budding event, both spatial and temporal aspects of bud emergence need to be considered. Here, we elucidate the complex interplay between the relative diffusion, the size of the pole, the time to polarisation and the concentration of active Cdc42 in the pole suggesting that perhaps cells have evolved multiple ways of maintaining this evolutionarily conserved phenomena.

## Methods

The representation of the parameter space (Fig 2) has been generated using Matlab [23]. For the simulations, a combination of an adaptive solver based *Finite Differences* (*FD*) in time and the *Finite Element Method* (*FEM*) in space was implemented (see section S2.2 in the Supplementary Materials for details). As the numerical implementation solves a system of PDEs, a spatial discretisation is required in terms of a *mesh* over the domain Ω (Fig 1A). For this purpose, the mesh was generated using the three-dimensional finite element mesh generator Gmsh [11]. For computational speed, we have implemented a non-uniform mesh with higher node-density close to the membrane and lower node-density in the cytosol as the former region requires more computational accuracy during the cell polarisation simulations than the latter. For the FD- and FEM-implementations, the computing platform FEniCS [2, 21] has been used. The visualisations (Fig 3, S4 and 6) have been constructed using the software ParaView [1, 3]. For all the simulations, we have used a cytosolic diffusion coefficient of *D* = 10, 000 and the spatially inhomogeneous initial conditions are perturbed around the steady states of the three states. To quantify polarisation properties, an empirical pole-recognition algorithm has been developed and implemented. A detailed description of the implementations can be found in the Supplementary Material (Supplementary Material S3).

The simulations have been conducted on two computational clusters. The first is a Dell PowerEdge R730 with an Intel Xeon E5-2683 CPU and an NVIDIA Tesla K80 GPU. The second computational cluster is based on an Intel Xeon Platinum 8180 CPU. The total simulation time of all simulations presented in the paper was approximately one week.

## Acknowledgements

This work was supported by the Swedish Foundation for Strategic Research (grant nr. IB13-0022 to MC and JB and grant no. AM13-0046 to PG) and by Vetenskapsrådet (grant no. 2014-6095 to AM). The authors would like to thank Martin Raum for aiding in the use of one of the two computational clusters used for the simulations and Axel Målqvist who discussed and provided references to the proof of theorem 1.

## Supplementary material

### S1 Analytical results

The analytical results consists of three parts: (1) A detailed description of as well as a motivation behind the bulk-surface activator-inhibitor system; (2) the details of the non-dimensionalisation procedure; (3) the proofs of all theorems 1, 2 and 3 on page 8 to 9 in the article.

#### S1.1 Description of the bulk-surface Activator-Inhibitor system

##### Influx of Inactive Cdc42 from the cytosol

The influx of inactive Cdc42 to the vicinity of the membrane, is determined by the concentration of GDI-bound Cdc42 in the cytosol (Fig 1C). The influx of the inactive GDP-bound form of Cdc42 is proportional to *G* where the rate of the reaction is determined by the rate constant *k*_1_ with units m^3^*/*min. We also assume that there exists a saturation level of active and inactive form of Cdc42 on the membrane, denoted by *k*_max_. These two assumptions result in the reaction rate given by:

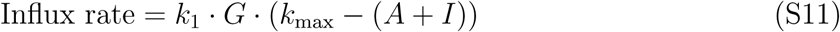

##### Dissociation of Inactive Cdc42 from the membrane

For the dissociation of GDP-bound Cdc42 from the membrane, we assume a first order reaction with rate constant *k*_−1_ with units m*/*min, which results in:

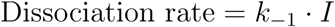

##### Activation of Inactive Cdc42

The activation of inactive Cdc42 which occurs on the membrane corresponds to the conversion of GDP-bound Cdc42 to the GTP-bound form. The reaction rate is assumed to be proportional to the concentration of active, GDP-bound form of Cdc42, and the rate constant *k*_2_ with units min^−1^ to be proportional to the concentration of GEFs, which we take to be constant during the time scale on which polarisation occurs. This results in a first order reaction term given by:

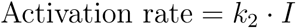

##### Inactivation of Active Cdc42

The inactivation of active Cdc42 corresponds to the conversion of GTP-bound Cdc42 to the GDP-bound form. Similar to the activation rate, it is also assumed to be a first order reaction, where the rate coefficient *k*_−2_ with units min^−1^ is assumed to be proportional to the concentration of GAPs, taken to be constant on the time scale of polarisation. This results in the following reaction rate:

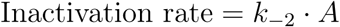

##### Activation of Cdc42 through a positive feedback loop

The feedback loop consists of the binding of active Cdc42 to PAKs which forms a complex which can further bind to various scaffolds. Together, this sequence of events forms a structure which can bind more GEFs and thereby enhance the activation process. During the time scale that polarisation occurs, we assume that both the concentration of PAKs and GEFs are constant, and that the feedback loop that recruits GEF is nonlinear. Such a feedback mechanism has been proposed previously [S13] and takes the form:

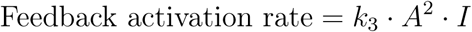

where the reaction rate coefficient is *k*_3_*A*, and requires both *A* and *I* to occur. The unit of *k*_3_ is m^4^*/*min.

#### S1.2 Non-dimensionalisation of the model

The aim to render the following model

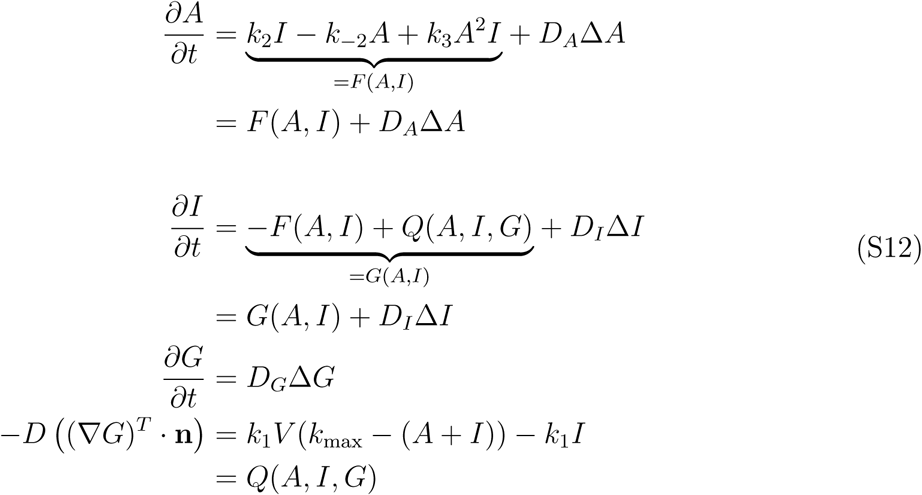

dimensionless. To this end, we follow the standard procedure in the setting of dynamic models in biology, and non-dimensionalise the model in order to reduce the number of parameters. Our choice of non-dimensional parameters are similar to the ones in the classical Schnackenberg model [S35]. Firstly, we introduce the following dimensionless states

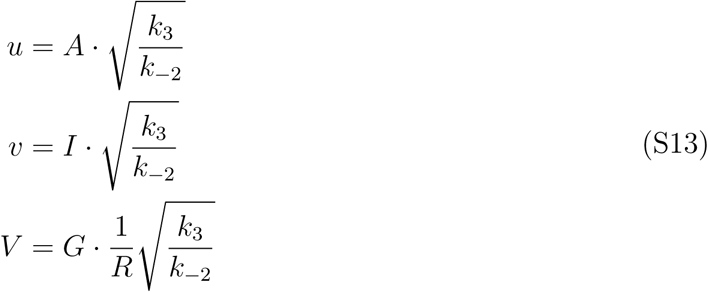

and the following dimensionless variables

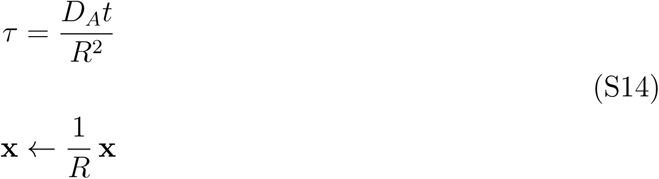

Note, that after the introduction of the spatial variable **x**, the domain Ω is transformed to the *unit ball* 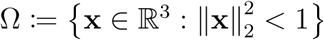 where the membrane described by 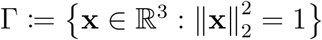 corresponds to the *unit sphere*.

By substituting the proposed scalings of the states in (S13) and the variables in (S14) we will derive the non-dimensional version of the model in (S12). We start with the left hand side and the time derivatives in order to obtain,

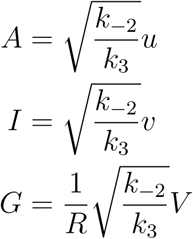

where the time variable is the following

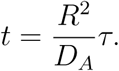

These expressions result in the following left hand side in the PDEs (S12)

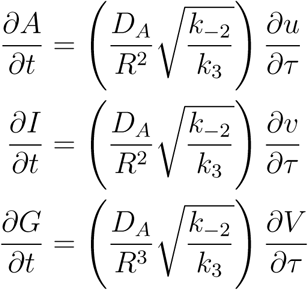

and thus the aim is to factor out 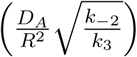 from the remaining terms in (S12): *F* (*A, I*), *G*(*A, I*), and the diffusion terms. For the diffusion terms, in the one-dimensional case^S1^, i.e. *x* ∈ [0, *L*], it follows that the following holds

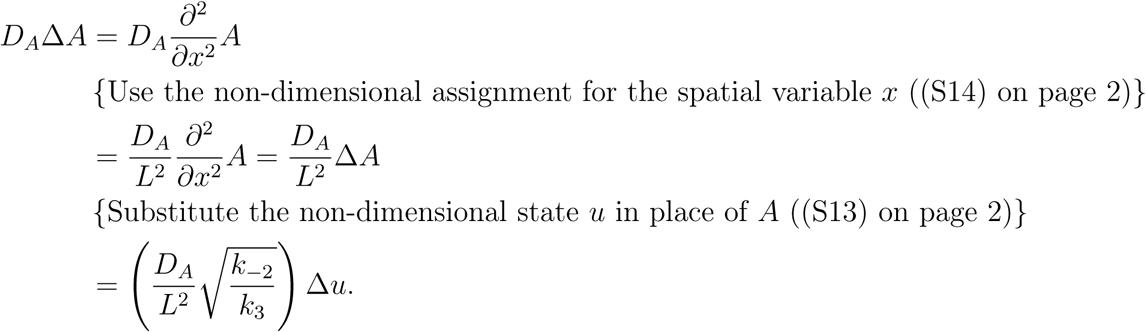

Analogously, in the spherical case with the spherical Laplace operator the factor *L*^2^ in the denominator in the one-dimensional case is replaced by the square of the radius *R*^2^ which is summarised as follows

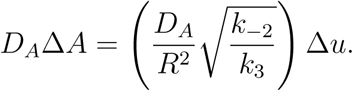

Similarly, for *I*

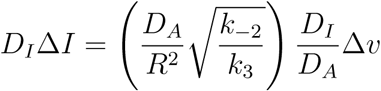

and for *G* the following holds

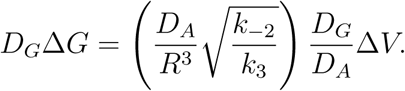

For the reaction term *F* (*A, I*), we have

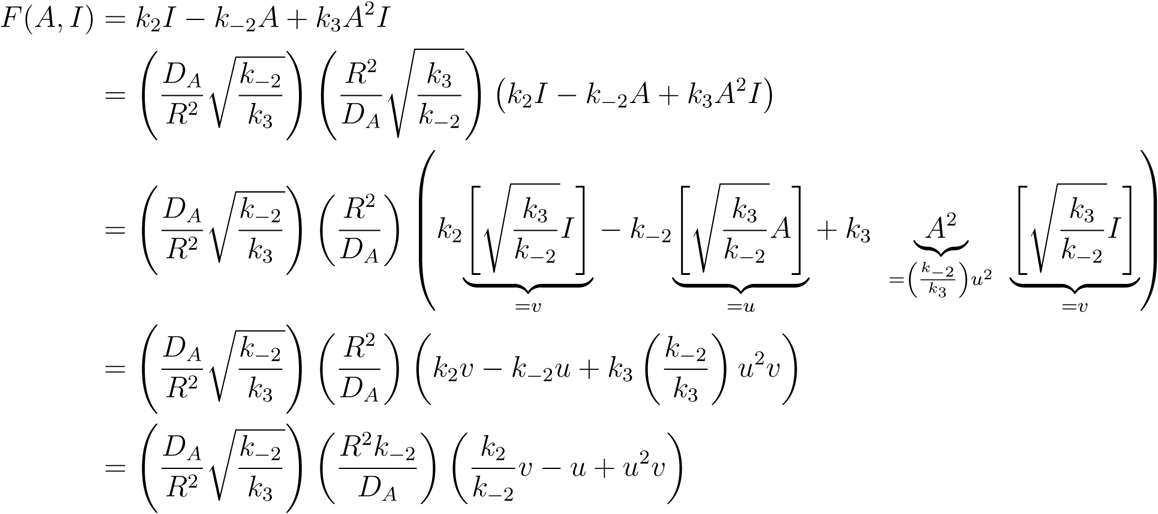

which is summarised as follows

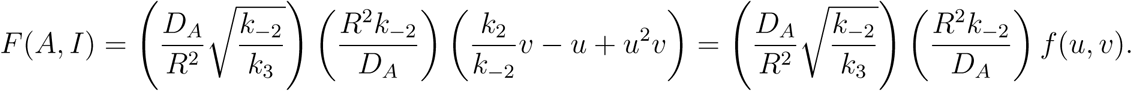

Lastly, we would like to factor out 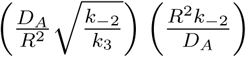 from the above expression of *F* (*A, I*) from *G*(*A, I*). However, as *G*(*A, I*) = − *F* (*A, I*) + *Q*(*A, I, G*) it suffices to look at the the transfer function *Q*(*A, I, G*). In a similar fashion, we obtain

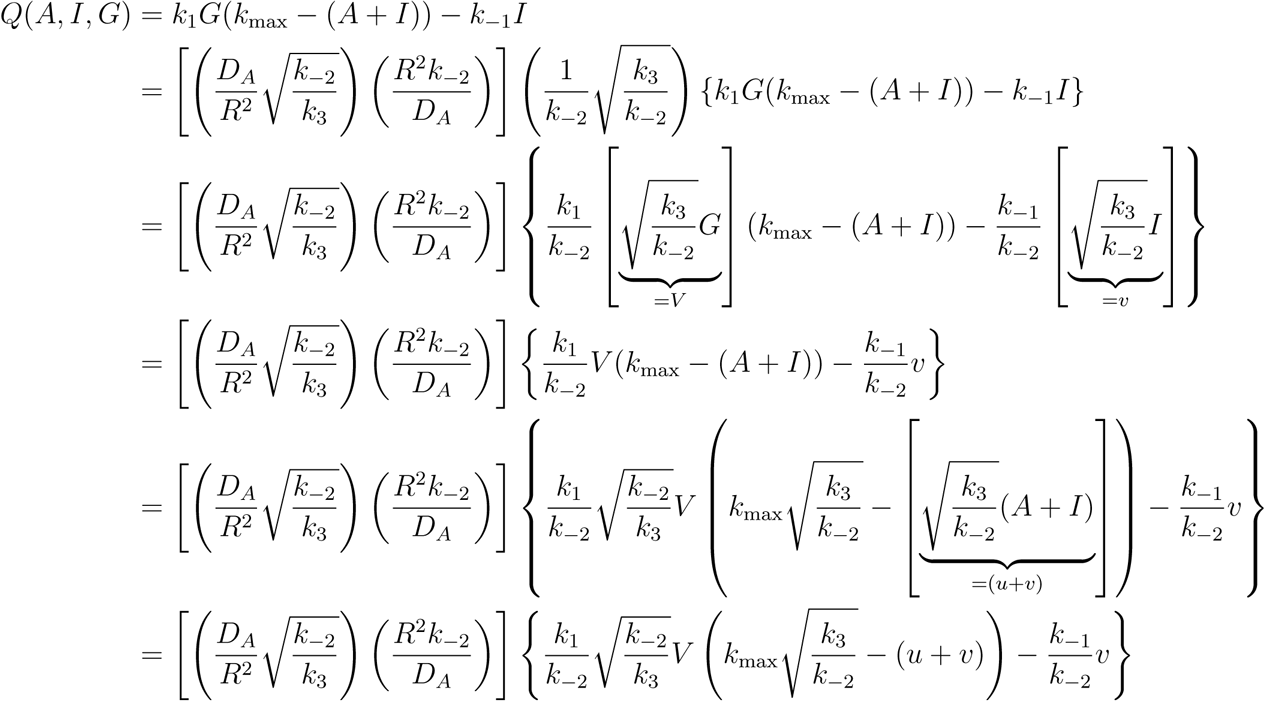

which yields the following result

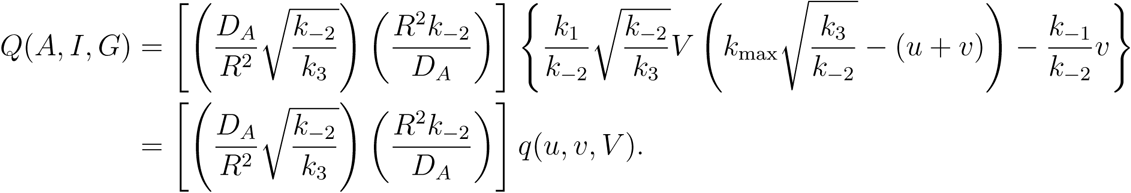

Now, summarising all these terms yields the following

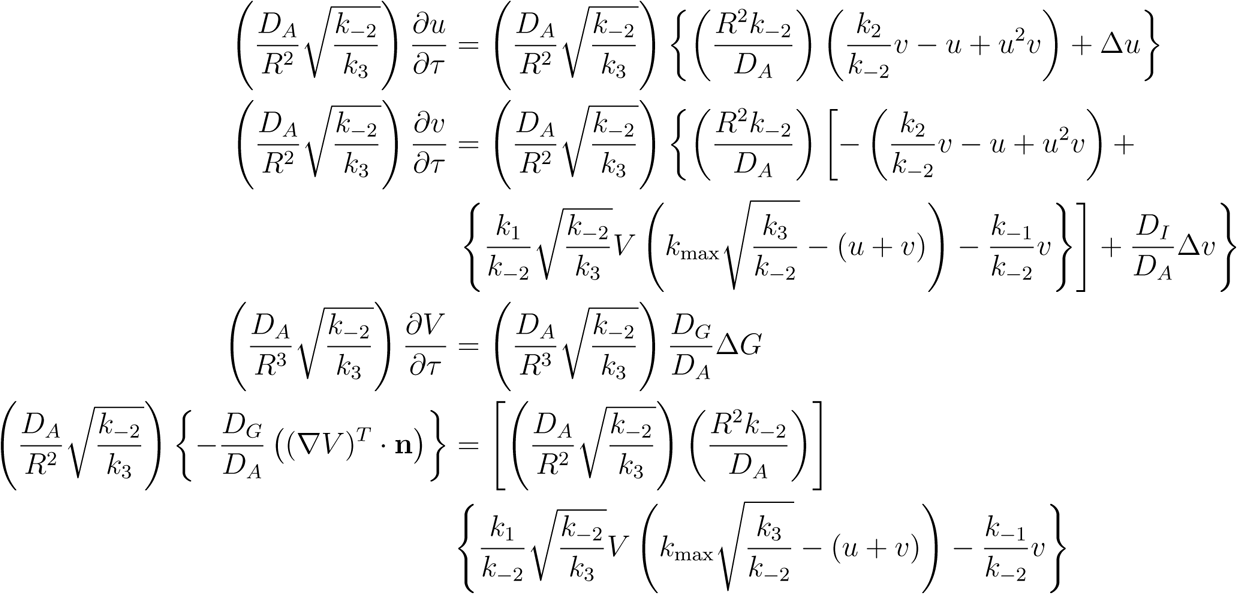

and cancelling the common factor in both sides results in the following equation

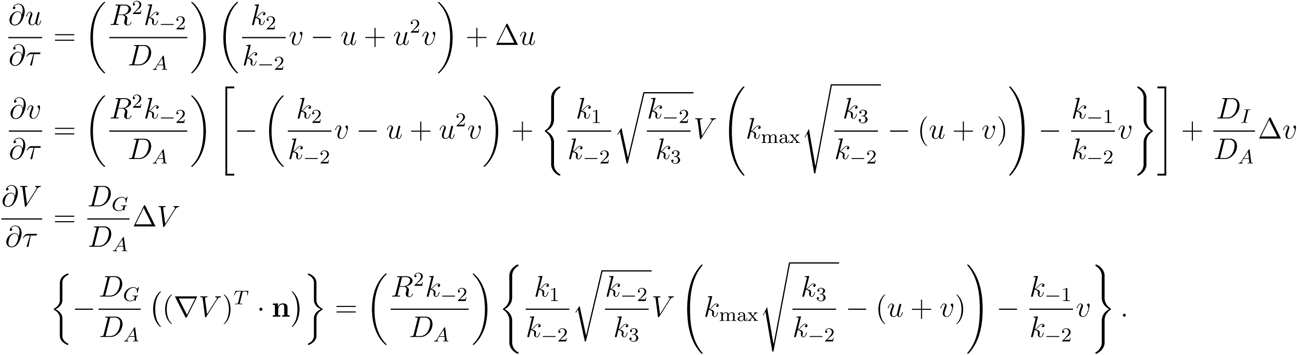

The introduction of the following parameters

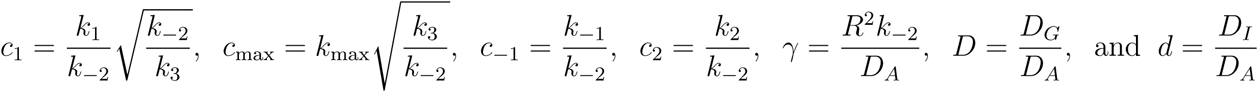

results in the following equations

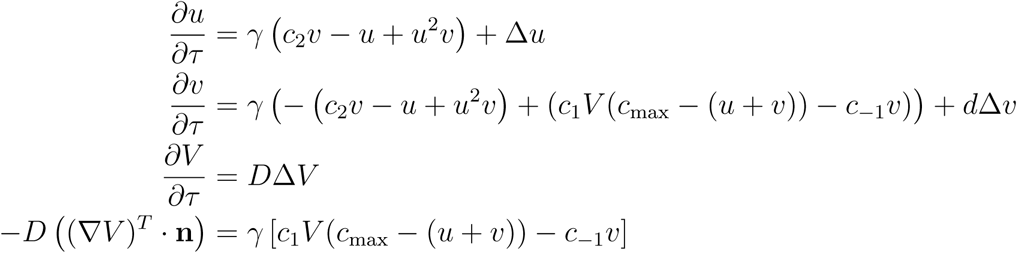

which is the desired result.

□

#### S1.3 Proof of Theorem 1

The proof is given by re-writing the PDEs on integral form using Duhamel’s principle and then use Banach’s contraction theorem on the corresponding mappings resulting from these integral forms. Similar existence proofs are well-studied for other RD models, see for example [22], but in this case the proof must be adapted for the unit sphere.

To this end, we equip the manifold Γ with a measure d*ω* defined by

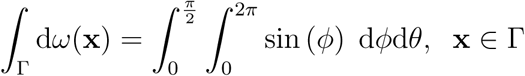

which implies that standard spherical coordinates corresponding to the angle *θ* ∈ [0, 2*π*] and the angle *ϕ* ∈ [0, *π*] are implemented. Also, we define the corresponding *L*_2_-inner product and norm as follows

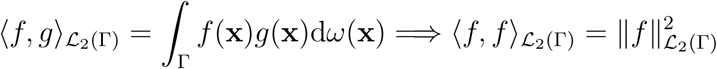

where we define the related ℒ_2_-space as follows

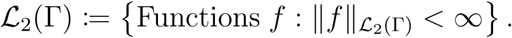

Using this norm, we can also define the ℋ^1^(Γ)-norm according to

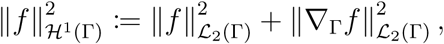

where ∇_Γ_ is the surface gradient in a weak sense with respect to Γ. This so called ℋ ^1^-norm requires that both the function value and its derivative do not “blow up”. Now, we can define our Sobolev space of interest, denoted by ℋ ^1^(Γ), as follows

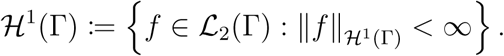

An *orthonormal* (*ON*)-basis for the Hilbert space of *L*_2_-functions on the sphere with the *L*_2_-scalar product defined above is the *Legendre polynomials* [S11] which we will denote by *P*_*n*_^S2^. Based on this, we can define two so called *solution operators* [S27] *E*_1_(*τ*) and *E*_2_(*τ*) respectively acting on a function *v* as follows

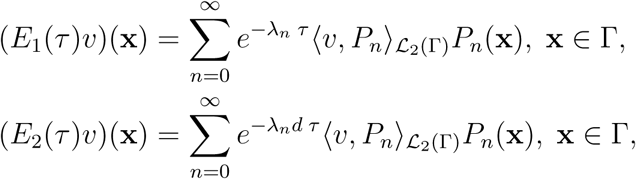

where the solution operator corresponds to calculating the Fourier series a given function *v* ∈ ℒ _2_(Γ). Here, the parameters *λ*_*n*_ correspond to the eigenvalues of the Laplace-Beltrami operator, and for an asymptotic expansion of the various eigenvalues *λ*_*n*_, see [S18]. Moreover, by using the inequality “*xe*^−*x*^ ≤*C*” for the exponential function it is possible to show the following bounds

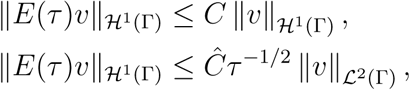

for a general solution operator *E* (i.e. the above bounds hold both for *E*_1_ and *E*_2_). Given these solution operators, for a time *τ* ∈ [0, *τ*_max_], the solution components *u*(*τ*), *v*(*τ*) ∈ ℋ^1^(Γ) of (S19) can be written on integral form according to Duhamel’s principle

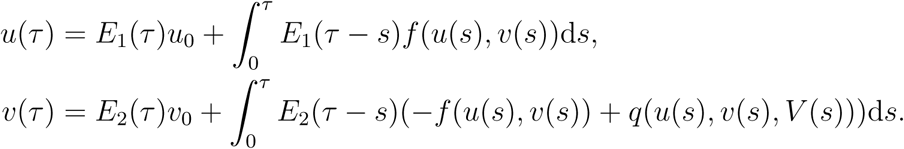

In fact, we can view these integral forms as operators *T*_1_ and *T*_2_ defined according to

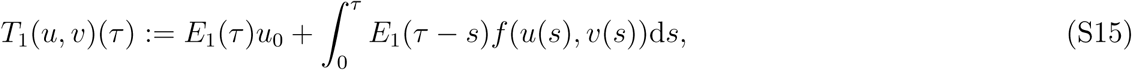

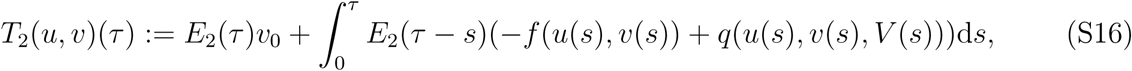

which strongly motivates us to consider the system on vector form. To this end, we denote the states on vector form as follows

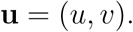

We define the operator **T** by, for a time *τ* ∈ [0, *τ*_max_]

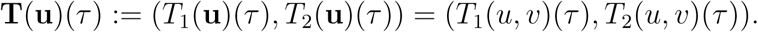

We note that for a time *τ* ∈ [0, *τ*_max_], **u**(*τ*) = (*u*(*τ*), *v*(*τ*)) ∈ ℋ^1^(Γ) × ℋ^1^(Γ). We equip this product space with the norm 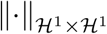 defined by

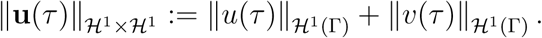

Let *X* = *C* ([0, *τ*_max_]; ℋ ^1^(Γ) × ℋ ^1^(Γ)) denote the Banach space of functions that are continuous in time on the interval [0, *τ*_max_] and for a fixed *τ*-value takes values in ℋ^1^(Γ) × ℋ^1^(Γ) (a so called Bochner space). Here, this space is equipped with the following norm

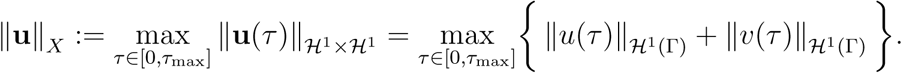

We now define a closed subspace of this Banach space by the ball ℬ as follows

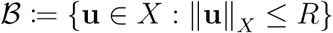

and furthermore we have the following bounds for the reaction functions *f, q*

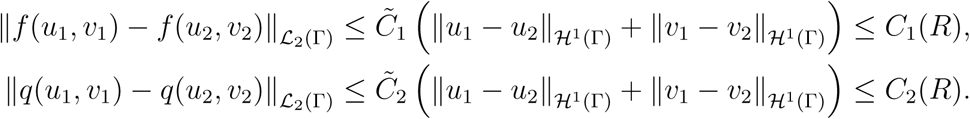

due to the fact that both these functions are continuous with continuous derivatives, i.e. *f, q* ∈ **C**^1^(ℝ^2^). In addition, since *f* (0, 0) = 0 and since *q*(0, 0) = *c*_1_*c*_max_*V*_0_ is bounded, we also have the following bounds

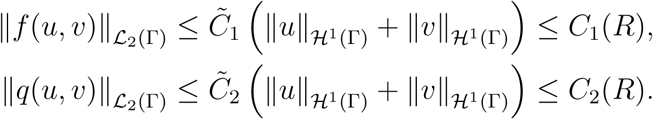

Note here that since the initial conditions *u*_0_, *v*_0_ are chosen in the region *A* the upper bound on these initial conditions are known, more precisely the following holds

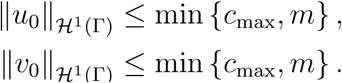

Next, we define a fixed point **u**^⋆^ ∈ ℬ as follows

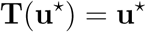

and the existence of a *unique* such point is guaranteed by *Banach’s fixed point theorem* [S10] if the following two conditions are satisfied:

1. The operator maps the closed subspace to the closed subspace, i.e. **T**: ℬ → ℬ. In other words, the image of the operator lies in the closed subspace of the Banach space, i.e.

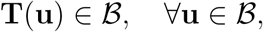
2. The operator is a contraction, i.e.

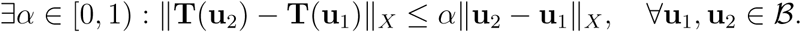

Regarding the first condition, we have

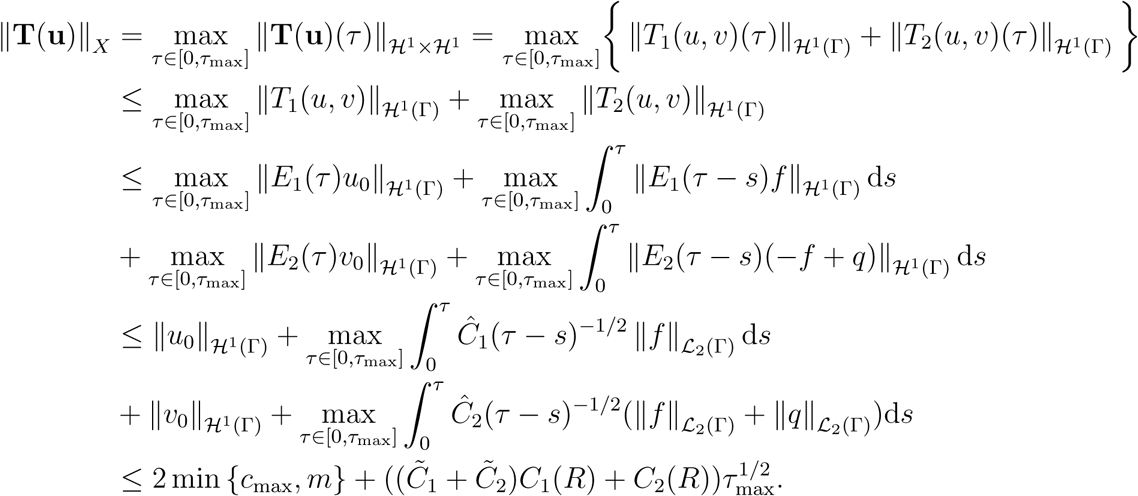

Now, if we choose

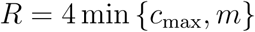

and by choosing *τ*_max_ sufficiently small so that

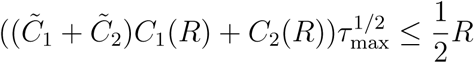

we have that ‖**T**(**u**)‖_*X*_ ≤ *R* and hence **T**: ℬ → ℬ.

In order to prove the second condition, we take **u**_1_ = (*u*_1_, *v*_1_) and **u**_2_ = (*u*_2_, *v*_2_) such that **u**_1_, **u**_2_ ∈ ℬ. It follows that

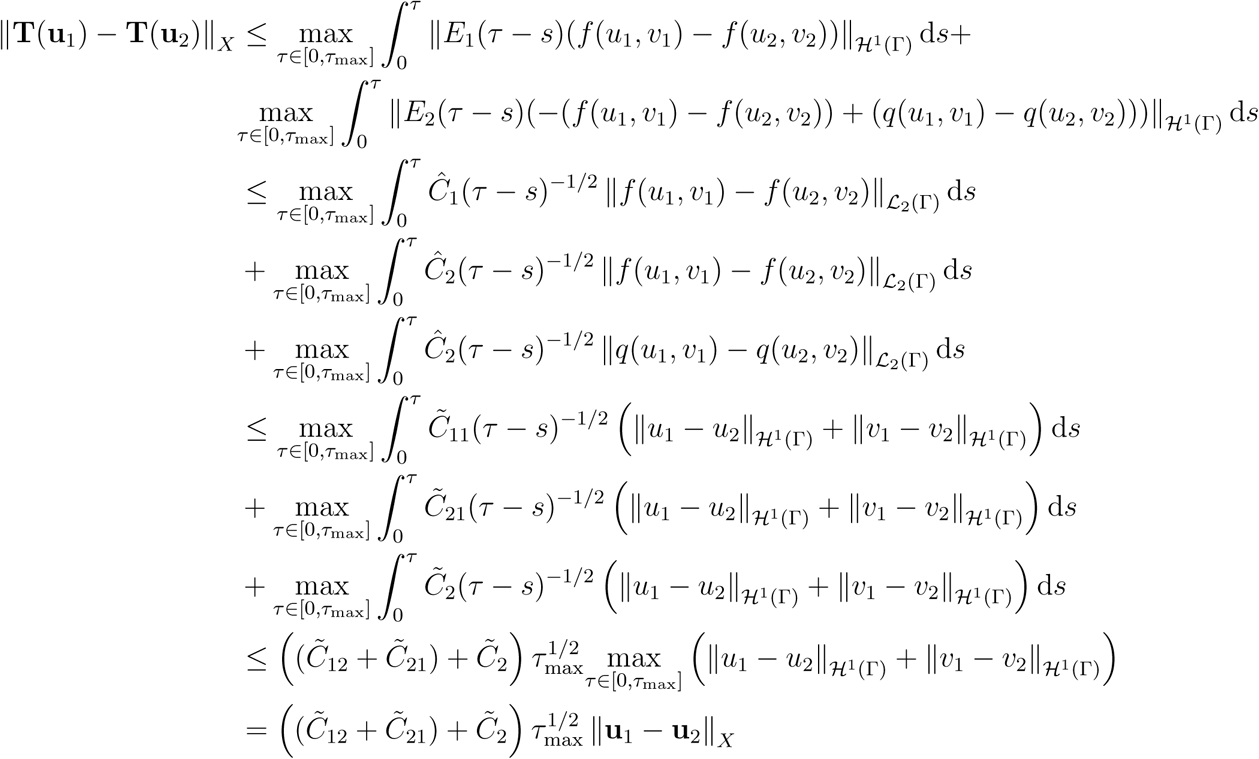

and thus choosing *τ*_max_ such that

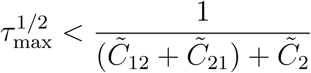

yields that **T** is a contraction. This shows we have a solution in local time, but in fact since the region *A* is a trapping region, it means that the solutions *u, v* are bounded in the ℋ^1^-norm for every *τ*, and from this follows that the existence of solutions is global in time.

#### S1.4 Proof of Theorem 2

The proof is based on the dynamics of the trajectories within the region *A* of the (*u, v*) state space defined as follows

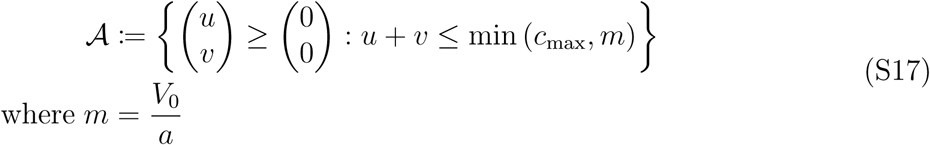

The same analysis for another choice of *f* has been conducted by Röger and Rätz in [S39]. To prove this we study the trajectories at the sides of this region. The region *A* is a trapping region if the following three conditions are satisfied for positive states *u, v >* 0:

1. ∂_*τ*_*u* = *f*(*u, v*) > 0 at *u* = 0,
2. ∂_*τ*_*v* = –*f*(*u, v*) + *q*(*u, v*) > 0 at *v* = 0,
3. ∂_*τ*_ (*u* + *v*) = *q*(*u, v*) < 0 at *u* + *v* = min{*c*_max_, *m*}.

For the homogeneous system, the functions *f* and *q* are defined as follows

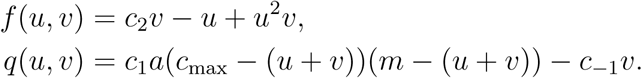

The first condition yields

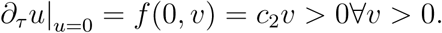

The second condition yields

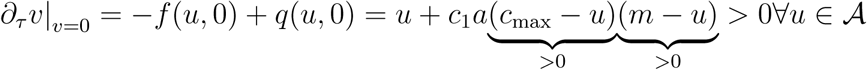

The third conditions, yields

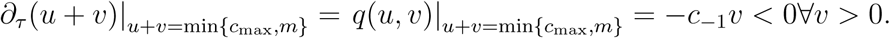

□

##### S1.4.1 Corollary 1 following from Theorem 2

The result follows directly from theorem 3.1 in [S38]. The trapping region above shows that the states satisfy the conditions known as “*positivity* “, i.e. *∂*_*τ*_ *v* | _*u*=0_, *∂*_*τ*_ *v*| _*u*=0_ *>* 0. Also, they need to satisfy what is called “*mass control*” meaning that *∂*_*τ*_ (*u* + *v*) should be bounded at all times. But this follows from the fact that *q* is bounded, i.e.

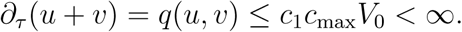

Also, the states are non-negative *u, v* ≥ 0 due to the positivity in combination with the positive initial conditions and we have a uniform upper bound on both states due to mass conservation.

□

#### S1.5 Diffusion driven instability in the limit *D* → ∞

We reduce the complexity of the system ((4) on page 5) by considering the limit *D*→ ∞. This implies that the cytosolic concentration of GDI-bound Cdc42 is approximated as being homogeneous motivated by the fact that the internal diffusion *D* is much faster compared to the diffusion in the membrane. In this case, the mass conservation property is described by the *non-local functional V* [*u* + *v*] below

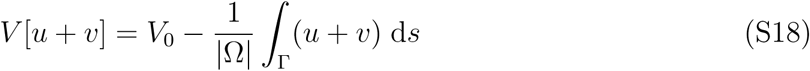

and the RD-system ((4) on page 5) gets reduced to the following two-state system.

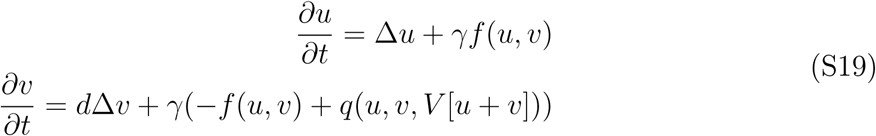

This system ((S18) and (S19)) was first described in Röger and Rätz [S40] a nd we w ill use the same notation as the one introduced in this work here. Now, the stability analysis concerns both the homogeneous system without spatial effects and the inhomogeneous system accounting for spatial effects. In the former case, the non-local functional is transformed to the non-local function *V*_1_(*u* + *v*) defined as follows.

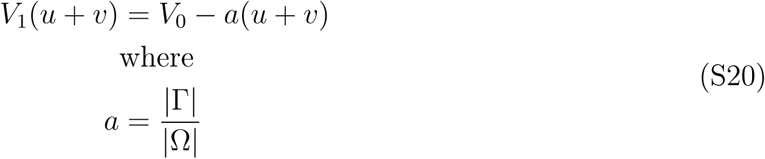

Note that in our case the domain Ω is the unit ball after the non-dimensionalisation and Γ is the unit sphere. Consequently, the parameter *a* (S20) has the value 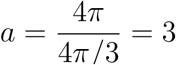.

Now, let (*u*^⋆^, *v*^⋆^, *V* ^⋆^) denote a steady state of the system (S19) and let *f*_*u*_, *f*_*v*_, *q*_*u*_, *q*_*v*_, *q*_*V*_ and 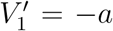 denote the partial derivatives evaluated at this steady state. Then, the stability of the homogeneous system is given by the following conditions.

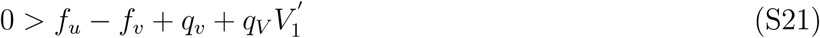

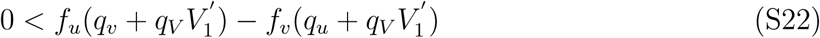

As in Proposition 3.1 by Röger and Rätz [S40], it is possible to obtain symmetry breaking in two ways provided that the above conditions ((S21) and (S22)) are satisfied. The first way is by the classic diffusion-driven instability proposed by Alan Turing [S44] corresponding to what we will refer to as the classic Turing conditions.

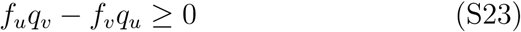

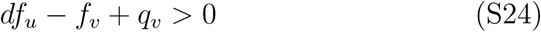

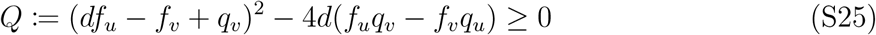

Here, the eigenvalues *λ*_±_ of the membrane bound Laplace operator Δ_Γ_ often referred to as the *wave numbers* are given by the following equation.

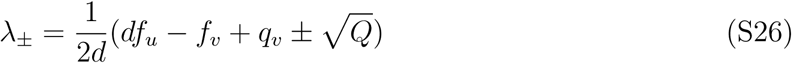

In fact, two conclusions can be drawn from the classic conditions ((S21) and (S24)). Firstly, the diffusion ratio must satisfy *d >* 1 implying that the state *v* diffuses faster than *u*. Secondly, if we denote the sign of the diagonal elements of the Jacobian matrix must be opposite. In linear stability analysis, the Jacobian matrix for a general system with reaction terms determined by *f* and *g* consists of the partial derivatives of *f* and *g* with respect to the states *u* and *v* where these derivatives are evaluated at the steady-state (*u*^∗^, *v*^∗^) of interest. As a consequence of the latter, it follows that the elements of the Jacobian matrix must have the following signs [S35, Fig 2.6 Page 88].

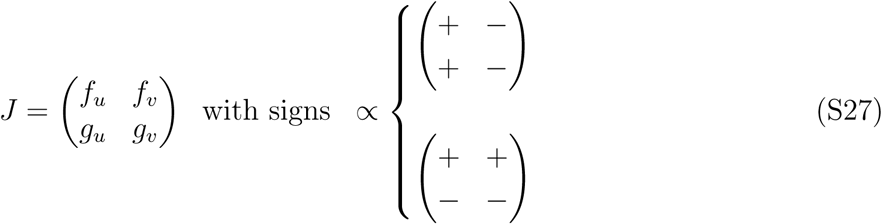

The other way by which symmetry breaking can be achieved which we will call *non-classic* Turing conditions are formulated as follows.

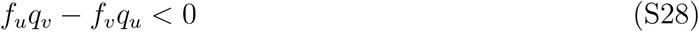

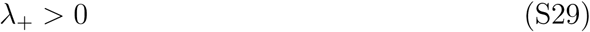

In the article, we state sufficient conditions for these two types of symmetry breaking by means of four theoretical results. The first lemma states conditions that are sufficient for a steady state that can give rise to Turing instability. The second lemma states conditions sufficient for a negative trace of the homogeneous system (S21). Given these two lemmas, the first theorem states sufficient conditions for Turing instability while the second theorem states sufficient conditions for saddle point instability. It is these conditions that we check numerically by firstly calculating a steady-state and then evaluating the conditions at the steady-state at hand. The details of how this is done will be presented in the next part of this document (S2.1). Before this is done, we prove Theorem 3 on page 9 stating the existence of steady-states and their characterisations.

#### S1.6 Proof of Theorem 3

We divide the proof of Theorem 3 into two parts. Firstly, we prove the number of steady states to the system. Secondly, we prove the existence of at least one steady state in the region *A* and then we derive a sufficient condition allowing for diffusion-driven instability.

##### S1.6.1 Existence of a steady state to the homogeneous system

Recall that a steady state is a solution (*u, v*) = (*u*^∗^, *v*^∗^) to the following two equations.

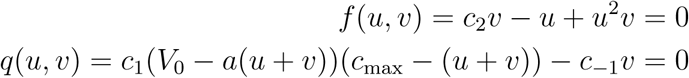

The first equation is satisfied for all *v* of the form

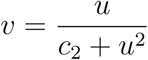

which can be inserted into the second equation, resulting in

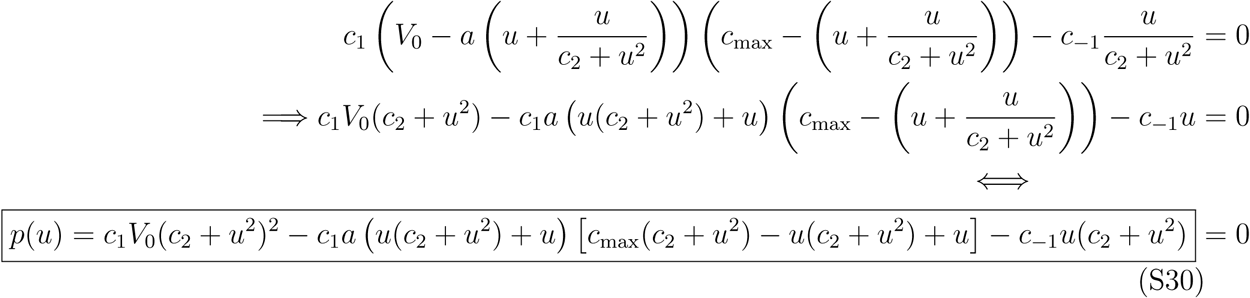

which is a 6^th^-degree polynomial in *u* denoted *p*(*u*). This polynomial can be simplified and written as follows.

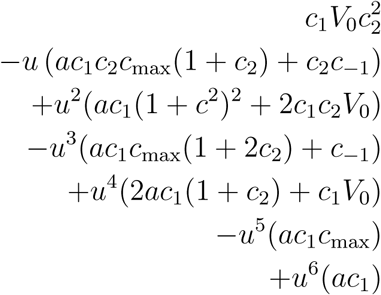

The solutions to *p*(*u*) = 0 (S30) subject to one additional condition (S17) which will be introduced in the subsequent section S1.6.2 are the steady-states of the original system.

Notice, that since all parameters in the expression above are positive, the coefficients of the polynomial are all positive in the case of even degree terms, and negative in the case of odd degree terms. We can therefore refer to Descartes’ rule of signs, to conclude that the polynomial has no negative real roots. We see this by observing that *p*(− *u*) has 0 sign changes between consecutive pairs of terms, and therefore there are no negative real roots. Being a 6^th^-degree polynomial, there can therefore be 0, 2, 4 or 6 positive real roots to the polynomial. However, it is worth emphasising that we are only interested in the roots of the polynomial *p* that satisfy a particular constraint as we mentioned earlier. This additional constraint states that an upper bound of the steady-states is 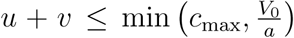, and therefore we analyse whether the system will have any steady-states satisfying this bound or not.

First, observe that 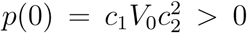. Next, we let *u*_1_ and *v*_1_ be such that *u*_1_ + *v*_1_ = min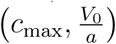 where *u*_1_ ≥ 0 and *v*_1_ ≥ 0. Given these values, we investigate

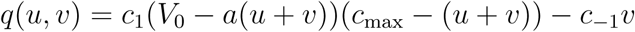

evaluated at *u* = *u*_1_, *v* = *v*_1_. We see that one of the two factors in the parentheses in the first term will be zero, and hence the whole expression is *q*(*u*_1_, *v*_1_) = *c*_1_*v*_1_ 0. Therefore, the sign of the polynomial at *u* = *u*_1_ will also be negative, i.e. *p*(*u*_1_) ≤ 0. By the intermediate value theorem, we can now deduce that at least once is *p* zero, inside the interval [0, *u*_1_].

##### S1.6.2 Characterisation of the steady-states enabling diffusion-driven instability

We seek a homogeneous steady state (*u*^∗^, *v*^∗^) ∈ *A* satisfying the following equations.

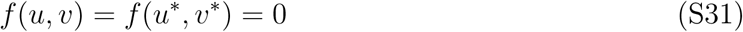

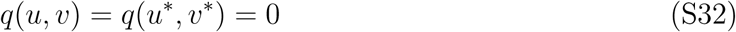

The first nullcline (S31) implies that the steady state of interest lies on the curve

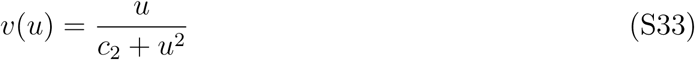

and thus we are interested in a *u*-component such that Φ(*u*):= *q*(*u, v*(*u*)) = 0. We next differentiate *v*(*u*) and find that

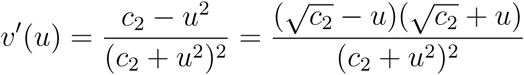

which has a positive root^S3^ at 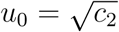, where 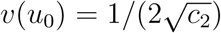. We also find that *v*′(*u*) *<* 0 for all *u > u*_0_ and that *v*′(*u*) *>* 0 for all *u < u*_0_. It is the switch of signs around the critical point *u*_0_ which we will use to characterise the desired steady-states.

Since, *f* (*u, v*) = *c*_2_*v* − *u* + *u*^2^*v*, the corresponding partial derivatives are

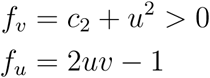

and since these terms are evaluated at the steady state we can substitute *v*(*u*) (S33) into the latter expression for *f*_*u*_ above to obtain the following.

**Figure S1:**
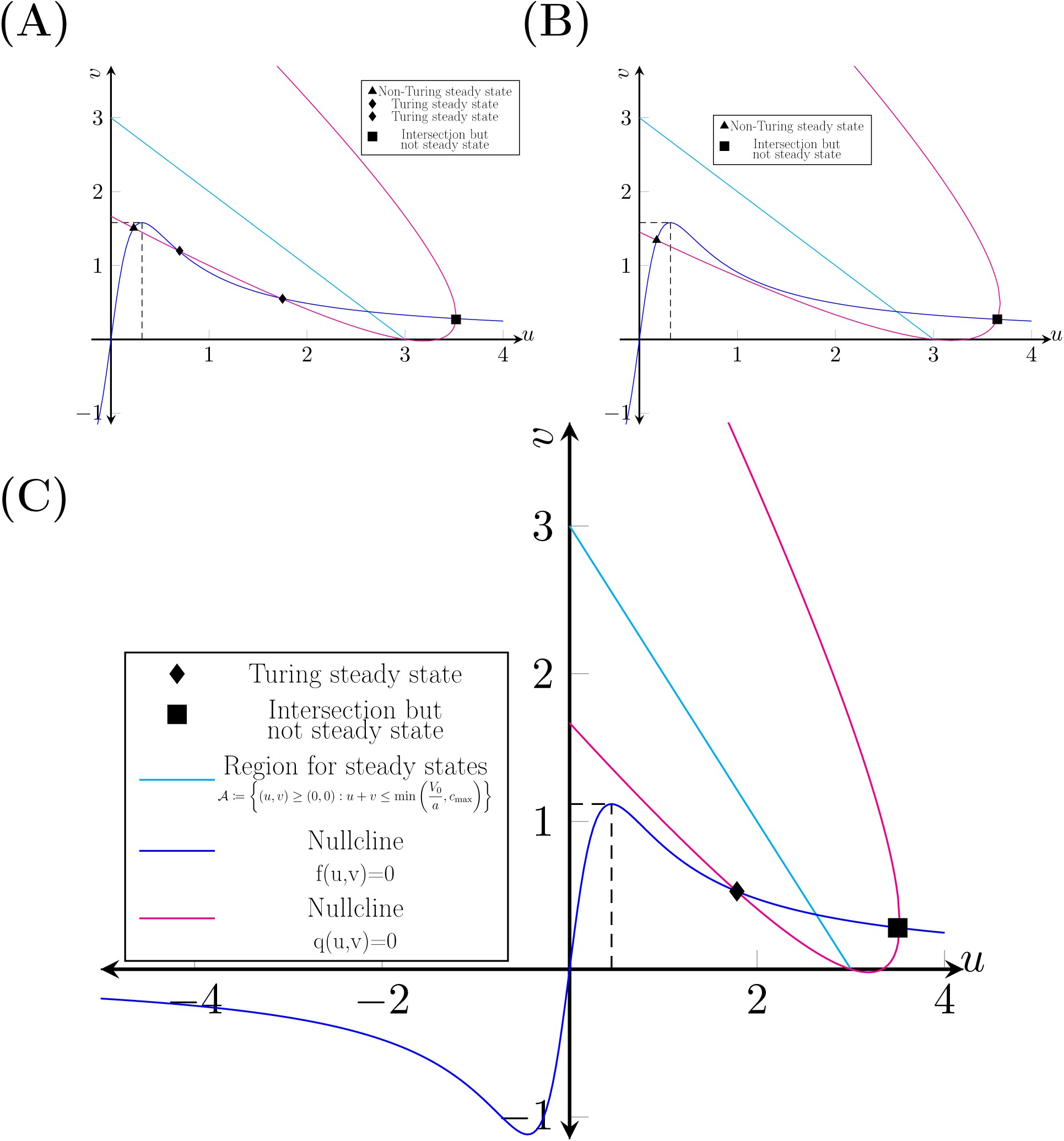
Steady States. The steady states of the system are located below within the region *A* in the first quadrant (the cyan curve). The intersections of two curves define the steady states: the nullcline corresponding to the activation-inactivation reaction *f* (*u, v*) = *c*_2_*v* − *u* + *u*^2^*v* (the blue curve) and the nullcline corresponding to the transfer of inactive Cdc42 between the cytosol and the membrane *q*(*u, v*) = *c*_1_ (*V*_0_ − *a*(*u* + *v*)) (*c*_max_ − (*u* + *v*)) − *c*_−1_*v* (the magenta curve). The number of steady states determined by the intersections of the two nullclines are visualised for three cases. **(A)** *Four steady states* (*c*_−1_, *c*_2_) = (0.30, 0.10): one non-Turing steady state ▴, two Turing steady states ♦ and one intersection but not a steady state ▪. **(B)** *Two steady states* (*c*_−1_, *c*_2_) = (0.30, 0.20): one non-Turing steady state ▴, and one intersection but not a steady state ▪. **(C)** *Two steady states* (*c*_−1_, *c*_2_) = (0.20, 0.20): one Turing steady state ♦, and one intersection but not a steady state ▪. The other parameters are: *V*_0_ = 10, *a* = 3, *c*_max_ = 3 and *c*_1_ = 0.05.

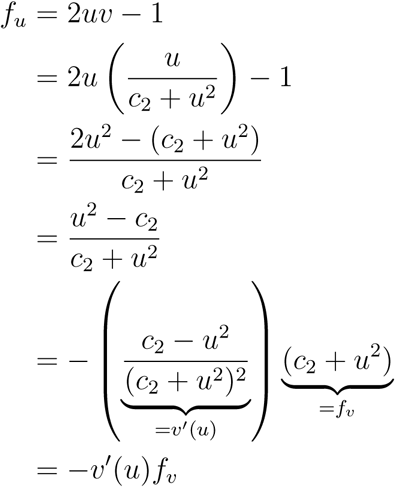

Now, the partial derivatives are evaluated at the steady-state (*u*^∗^, *v*^∗^) which can be characterised based on the sign of the derivative *v*_*u*_ at the steady-state. It is known that a steady-state allowing for diffusion-driven instability has a Jacobian matrix with elements with particular signs (S27). As a consequence of the negativity of the partial derivatives of *q*, the only way to obtain opposite signs of the diagonal elements *J* (1, 1) and *J* (2, 2) is if the following holds.

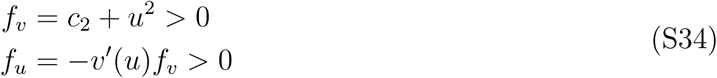

It is clear that *f*_*v*_ *>* 0 however *f*_*u*_ *>* 0 implies that *u*^∗^ *> u*_0_ for the critical point 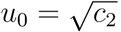. This follows from the fact that we have that *v*′(*u*) *<* 0∀ *u > u*_0_, and −*v*′(*u*) *>* 0 ∀ *u > u*_0_. Combining this with the fact that the steady-states should be located withing the region *A* (S17) yields the following characterisation of the desired steady-state.

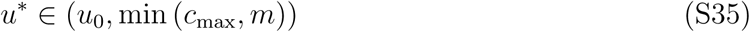

In fact, this requirement indicate that the signs of the Jacobian must satisfy the lower of the two cases in terms of the signs of the elements (S27). Also, a necessary requirement to ensure that such a steady-state exists is that the local maximum

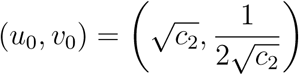

of *v*(*u*) (S33) is located within the region *A* (S17). This requirement is exactly translated to the condition

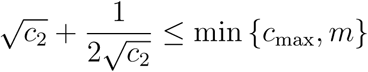

which completes the proof of Theorem 3.

### S2 Numerical implementations

This section consists of three parts: (**1**) A pseudo-code for the visualisation of the parameter space (Fig 2) giving rise to diffusion-driven instability, (**2**) The implementation of the FD-FEM based algorithm for solving the full system in (4) on page 5 and (**3**) We present an empirical pole-detection algorithm which terminates the simulation when the pole is formed.

#### S2.1 Numerical mapping of the parameter space

The numerical mapping of the parameter space consists of three steps. For the sake of simplicity, assume that the (*c*_−1_, *c*_2_)-space^S4^ is to be calculated. Then, the initial step is to discretise both the *c*_−1_- and *c*_2_-line into a finite number, *N* ∈ ℕ_+_, of nodes. Using this partitioning, allocate memory for the solution matrix 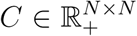, i.e. *C* ← **0**^*N*×*N*^. Then loop over the nodes in the partitioning and do the following two steps for all nodes.

1. Calculate the steady states (*u*^∗^, *v*^∗^) ∈ *A*
2. Check the desired conditions for diffusion-driven instability
  a. **Classic case**: Check the conditions (S21), (S22), (S23), (S24) and (S25). If they are satisfied, assign *C*(*i, j*) ← 1.0 for the specific indices *i, j* ∈ {1, *…, N* } of interest
  b. **Non-classic case**: Check the conditions (S21), (S22), (S28) and (S29). If they are satisfied, assign *C*(*i, j*) ← 0.5 for the specific indices *i, j* ∈ {1, *…, N* } of interest

Depending on the specific case, a matrix with the value 1.0 for all parameters giving rise to classic diffusion-driven instability is obtained or a corresponding matrix with the value 0.5 in the non-classic case.

To calculate the steady-states, we used a Newton-iteration with a given start guess. To this end, we take the function *v* (S33) corresponding to the nullcline *f* (*u, v*) = 0. Similarly, we rewrite the nullcline *q*(*u, v*) = 0 as a function of *u* in order to obtain the following two segments.

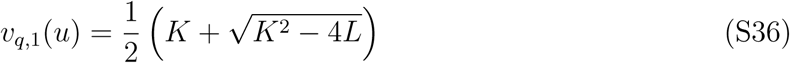

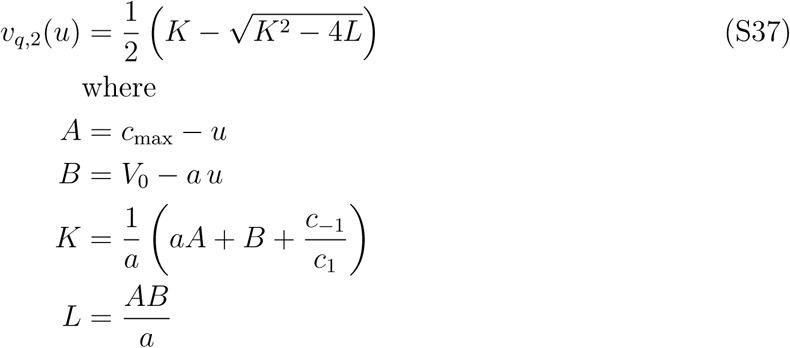

Now, given a start-guess *u* = *u*_0_, we solve the equations *v*(*u*) −*v*_*q*,1_(*u*) = 0 and *v*(*u*) −*v*_*q*,2_(*u*) = 0 numerically using the function fzero in Matlab [S34]. In order to make sure that the solver actually finds the steady-states, we take a large number of start guesses (in fact, we have used 50 start guesses) in the interval [*u*_0_, min (*c*_max_, *m*)] and we control that each value *u*^∗^ that the solver converges to satisfies the equation 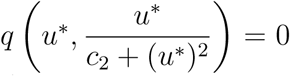. If so, the value (*u*^∗^, *v*(*u*^∗^)) is a steady state and it is consequently saved.

Lastly, given the steady state we can check the conditions involving the partial derivatives. The partial derivatives for our model are the following.

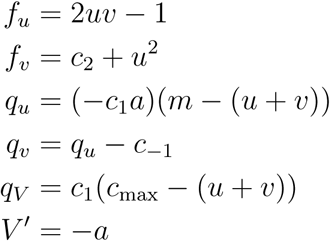

#### S2.2 Numerical solutions to the RD-model of cell polarisation

The numerical implementation of the solution to the problem is divided into three parts. Firstly, we set up the problem by formulating the equations of the model and the corresponding domains to the various equations. It is worth emphasising that in the analysis, it is assumed that the cytosolic diffusion goes to infinity, i.e. *D* → ∞while this is not the case for the numerical solutions. Moreover, as the variables in the PDE-problem are **x** ∈ ℝ^3^ corresponding to the *spatial* dimension and *t* ∈ ℝ_+_ corresponding to time, two discretisations corresponding to these variables are required. Firstly, a *Finite Element Method* (*FEM*) is implemented corresponding to the spatial discretisation. Secondly, a *Finite Difference* (*FD*)-method is implemented corresponding to the discretisation in time. These two discretisations are presented subsequently after the numerical setting is introduced.

##### S2.2.1 Introduction to the numerical implementation: Setting up the problem

The numerical solution to the original RD problem (Eq (4) on page 5) accounts for the domains for the problem (Fig S2). These are the cytosol Ω corresponding to the interior of the spherical cell and the cell-membrane Γ being its surface (Fig S2).

The domain (S38) and interfaces (S39) are important to define as the various equations and boundary conditions are based on the spatial description. Note that the domains are the unit sphere and its surface which correspond to the geometric description of the dimensionless model.

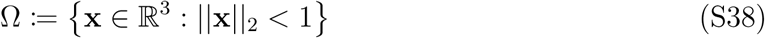

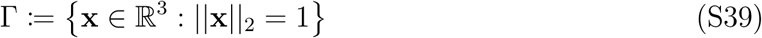

Now, given the definition of the spatial domain Ω the formulation of the problem is the following. In the membrane Γ the active *u* and inactive *v* form undergo activation-inactivation reactions determined by the function *f* (*u, v*) = *c*_2_*v* −*u* + *u*^2^*v* and diffusion. The interface Γ has a flux of the inactive membrane bound component *v* from the membrane to the cytosol, a flux of the GDI-bound cytosolic component *V* from the cytosol to the membrane and no flux of the active membrane bound component *u*. The flux over the membrane is determined by the function *q*(*u, v, V*) = *c*_1_*V* (*c*_max_ − (*u*+*v*)) −*c*_1_*v* and it is translated into a contribution to the reaction term for the inactive membrane-bound species *v* and a Robin-boundary condition for the cytosolic component *V* at Γ. Lastly, in the cytosol, Ω, the GDI-bound component *V* undergoes diffusion. Using these equations and boundary conditions it is possible to formulate the detailed model of Cdc42 activation (S40).

**Figure S2:**
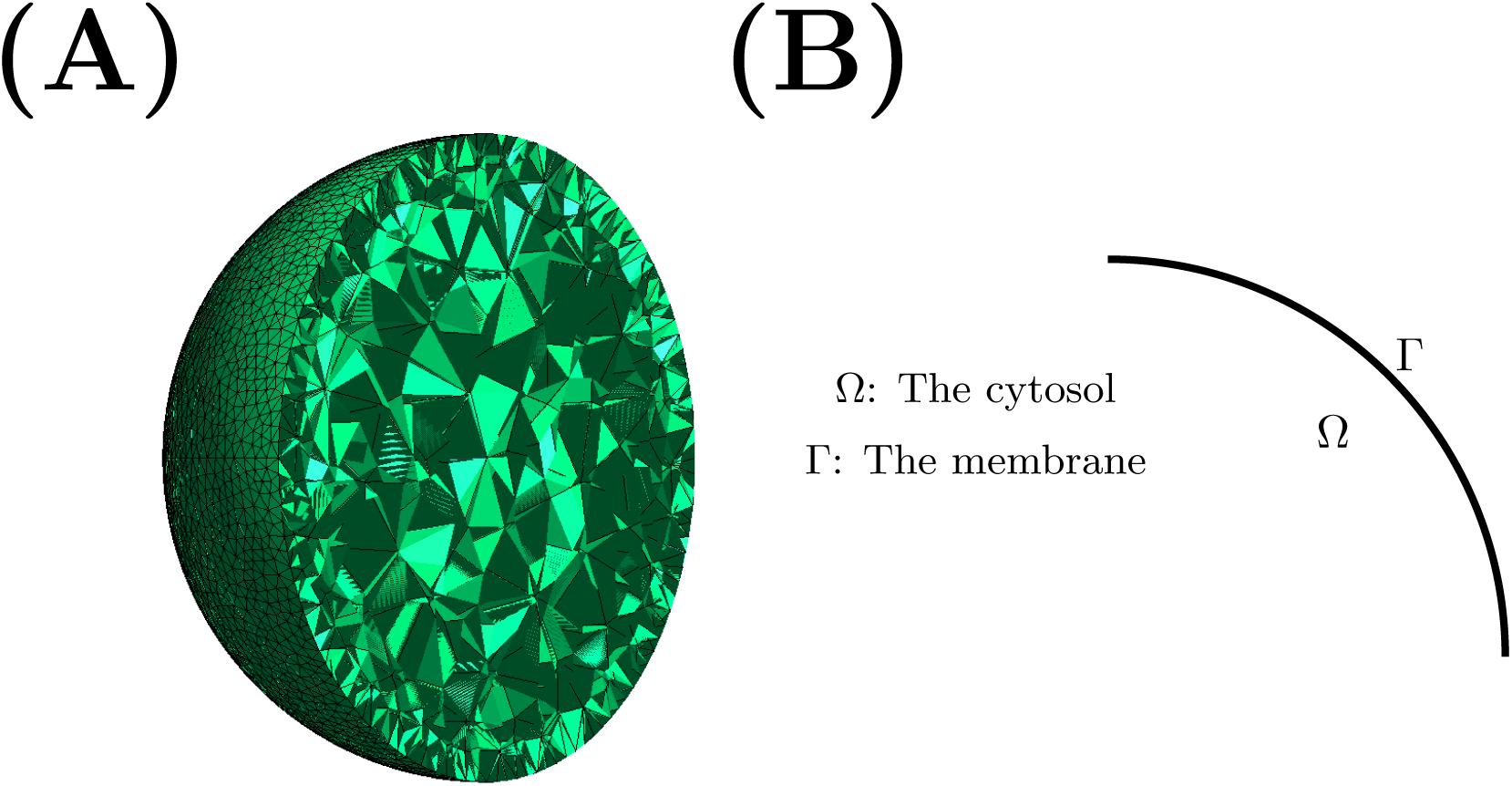
The detailed Spatial Domain. (**A**) *Spatial mesh for the numerical solution*. The discretisation of the spatial domain represented as a mesh. The mesh is non-uniformly discretised in the sense that the node-density is higher at the membrane than in the cytosol. (**B**) *Geometric domain*. By letting the membrane thickness shrink to zero, the geometric description is simplified to one domain Ω corresponding to the cytosol and one boundary Γ corresponding to the membrane

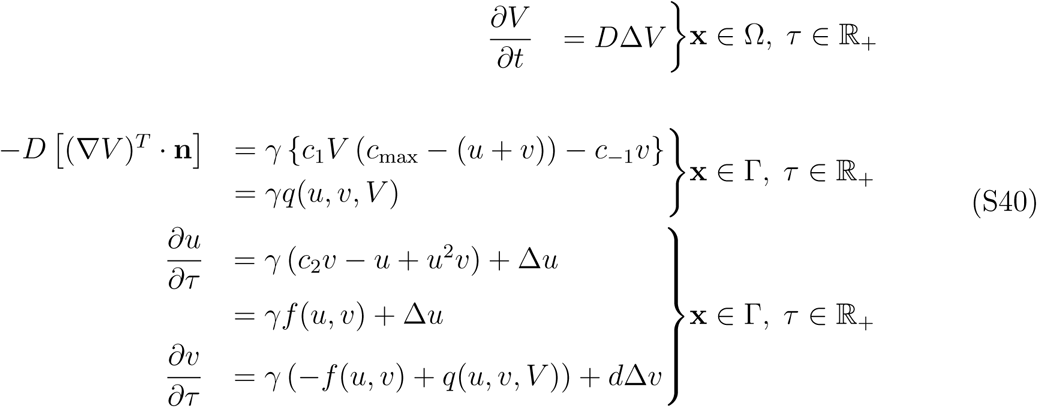

The various notations above are the following: 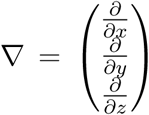 is the gradient operator, “*T*” is the transpose operator, **n** ∈ Ω ⊂ ℝ^3^ is the *outward normal* at a specific position on a given surface and 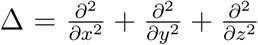 is the Laplace operator. In order to solve the problem (S40) numerically, the spatial domain must be discretised.

The spatial discretisation of the domain is corresponds to a *mesh* (Fig S2A). As the interesting parts of the model regards the reactions and diffusion of the membrane-bound species *u* and *v* it is advantageous from a computational view to use a non-uniform mesh. More precisely, we have implemented a mesh with higher node density close to the membrane and a low node density in the interior of the cell. To generate the mesh over the domain, we have used the three dimensional finite element mesh generator Gmsh [S12]. Given the generated mesh, it is possible to numerically solve the problem (S40) using the finite element method.

For the FD- and FEM-implementations, the computing platform FEniCS [S19, S21, S22, S25, S26, S23, S28, S36, S24, S29, S42, S4, S6, S32, S37, S14, S17, S41, S5, S2, S30, S33, S31, S20, S7, S3, S1, S15, S16] has been used. In FEniCS, it suffices to provide the so called *variational formulation* of the problem (S40) which is a reformulation of the problem. The approximation to this problem is then obtained by projecting the variational formulation onto a space of piece wise continuous functions or in our case a test function space of piece-wise linear functions. In the next chapter, we will merely present the derivation of the variational formulation regarding the spatial discretisation, and after that we will present the finite difference implementation for the temporal discretisation based on this variational formulation.

##### S2.2.2 An implementation of the Finite Element Method in space

Define the following function space for *test functions φ* on the domain Ω.

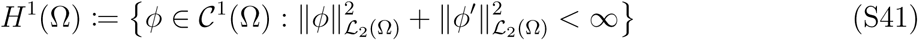

This space consists of continuous functions with continuous first derivatives with compact support.

As is standard in the formulation of the variational formulation, we multiply the three PDEs (S40) with three test functions *ϕ*_1_, *ϕ*_2_, *ϕ*_3_ ∈ *H*^1^(Ω) and integrate over the domain, Ω.

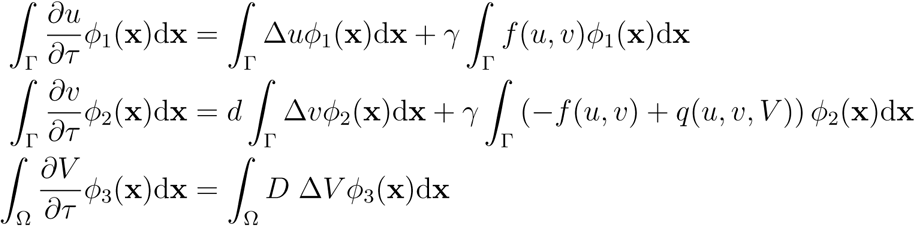

Now, to rewrite the diffusion terms, we will use *Green’s first identity* [S27]

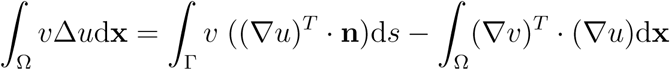

where 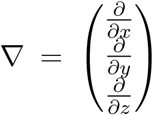 is the gradient, **n** ∈ ℝ^3^ is the outward normal and ^*T*^ is the transpose operator.

Using this identity on the diffusion terms above one obtains

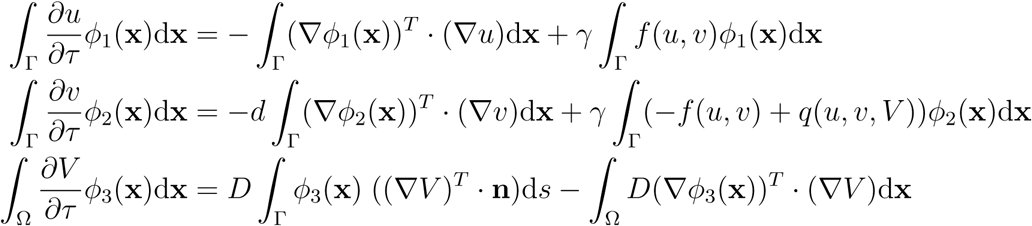

and simplifying the above yields the following.

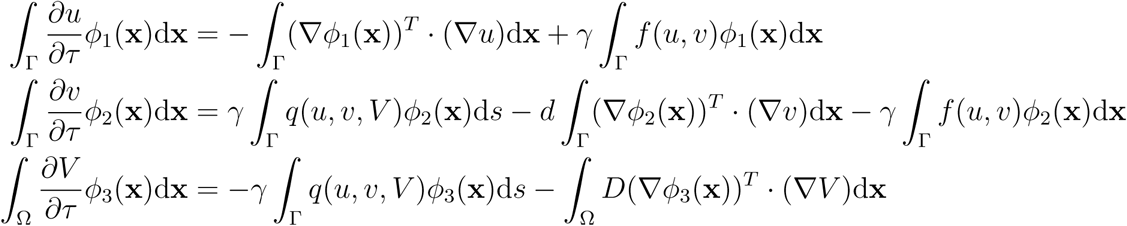

Lastly, moving all terms in the right hand side to the left hand side and adding all three equations yields the *variational formulation* of the detailed problem (S40).

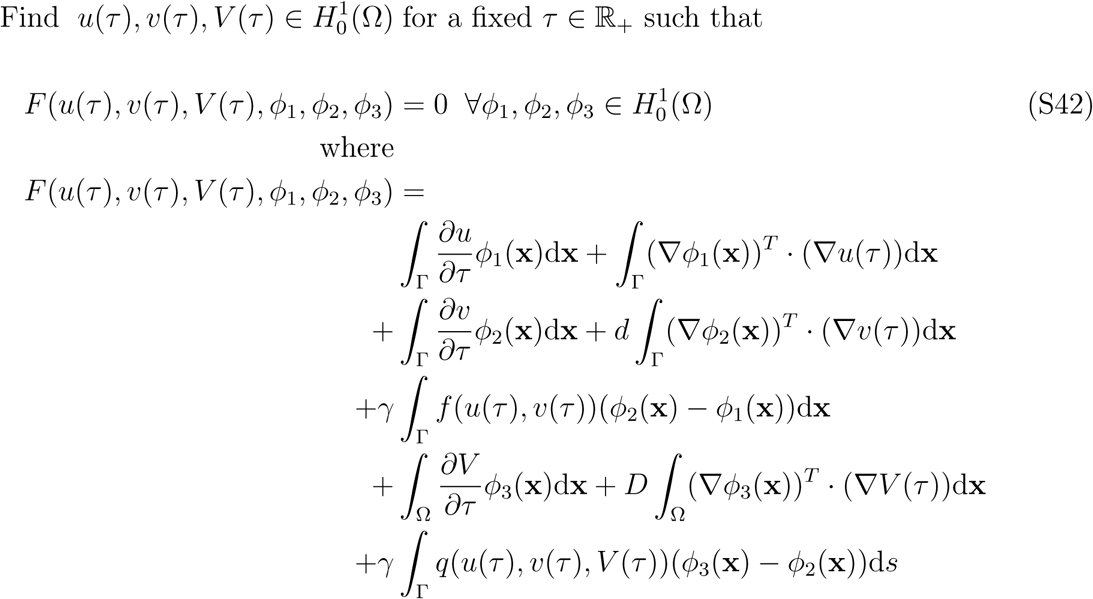

The numerical solution of this problem (S42) is written as linear combinations of piece-wise continuous basis functions of order 1, i.e. linear functions. Assume that a spatial discretisation (Fig S2A) is given and denote this by *τ*_*h*_. To describe the finite element method, denote the space of piece-wise linear functions on *τ*_*h*_ by *ℌ* (Ω). These tetrahedron-like functions take the value one on each node in the grid (Fig S2A) and they take the value zero on the neighbouring nodes. Then, the finite-element solution *u* (*τ*), *b* (*τ*), *ቨ* (*τ*) ∈ *ℌ*(Ω) for a fixed *τ* ∈ ℝ_+_ to the variational formulation (S42) consists of the orthogonal projections of 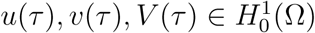 onto *ℌ*(Ω) for a fixed *τ* ∈ ℝ_+_. Here, we mean projection in the sense of the classic ℒ_2_*-inner product*.

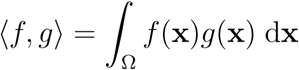

It is this variational formulation (S42) that is solved in FEniCS approximately using the finite element method just described. Subsequently, the time derivatives will be approximated using a finite difference scheme.

##### S2.2.3 An implementation of the Finite Difference Method in time

For the implementation of the finite difference scheme for solving the problem (S42) numerically, we use an *mixed Implicit-Explicit scheme*. The solution of the ODE problem

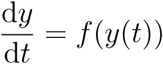

with *f* ∈ **C** (ℝ) is the function *y*∈ **C** (ℝ _+_) with the time as variable *t* ∈ ℝ _+_. Initially, the general methodology for finite differences involves *discretising* the time-line into various nodes, where the partitioning of the time-line is denoted as follows *T* ([0, *t*_max_]) = {*t*_0_, *t*_1_, *t*_2_, *…, t*_*n*_, *…, t*_max_}. Moreover, denote each of these nodes by *t*_*n*_ ∈ ℝ _+_ for some index *n* ∈ ℤ_+_ where the previous node on the discretised time line is denoted *t*_*n*−1_. Then, the backward-Euler algorithm for the above problem entails solving solve the following equation iteratively

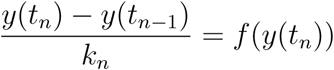

where *k*_*n*_ = *t*_*n*_ − *t*_*n*−1_ ∈ ℝ _+_ is the so called *step size*. The solution to the above equation corresponds to the solution *y*(*t*_*n*_) in the current node given the solution in the previous node *y*(*t*_*n*−1_). Note that both the left and the right hand side in the above equation depend on the solution *y*(*t*_*n*_) in the current node which implies that this algorithm is *implicit*. The forward version of the Euler algorithm consists of solving

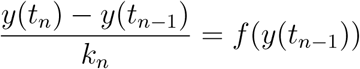

which yields the iterative scheme *y*(*t*_*n*_) = *y*(*t*_*n*−1_) + *k*_*n*_*f* (*y*(*t*_*n*−1_)) which is *explicit* and can be solved directly. However, the implicit version is more computationally expensive and has to be solved by using for example Newton’s method [S9, S8]. Despite the implicit algorithm being more computationally expensive compared to the forward or explicit version of the Euler scheme for ODEs, the advantage of it is that it is *unconditionally stable* as oppose the the explicit algorithm which is merely *conditionally stable*. The numerical solution of the detailed problem (S40) is given by combining the finite element method based on the previous variational formulation (S42) with a mixed explicit-implicit Euler finite difference scheme described above, which results in the following variational formulation (S43).

Find 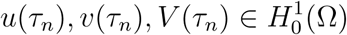

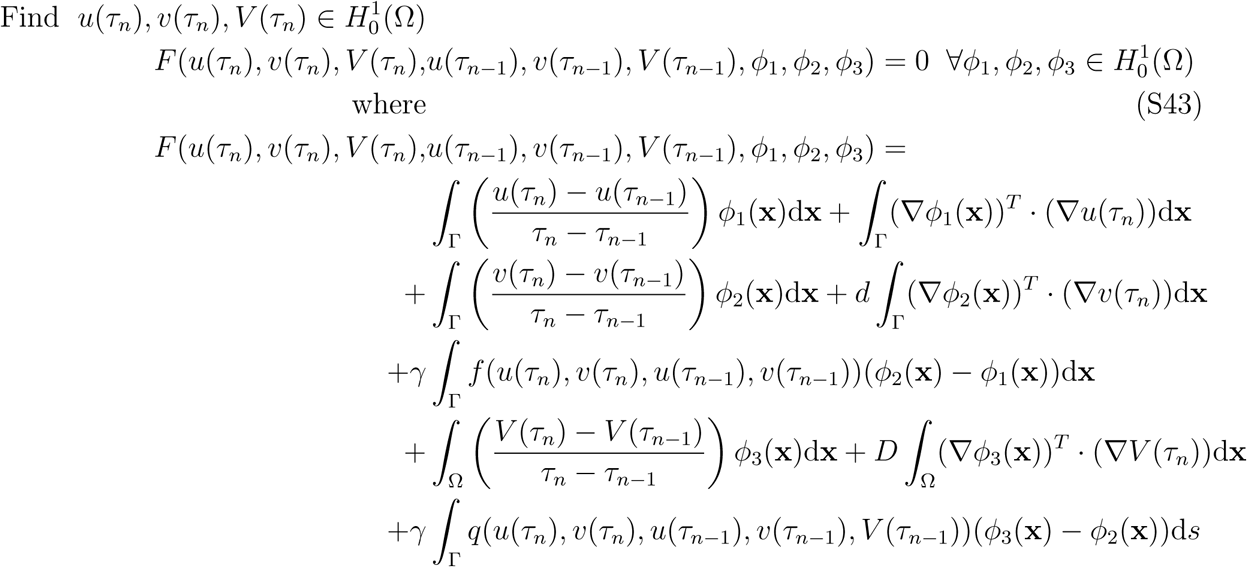

The above approach is referred to as an *mixed Implicit-Explicit* approach. This approach is implicit in the sense that the linear terms, i.e. the time derivatives approximated by finite differences and the diffusive terms containing the gradients, are implicit. These will result in the classic mass- and stiffness-matrices for the time derivatives and diffusive terms respectively when the finite element method is implemented. By “trial and error” we found that a *mixed implicit-explicit* methodology worked in order to get accurate simulations.

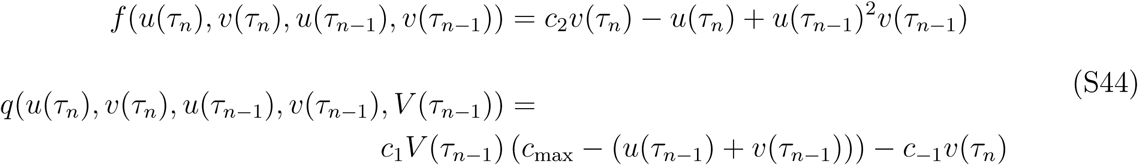

As can be seen, all purely linear terms are treated implicitly while the purely non-linear terms are treated explicitly. Furthermore, we implemented an adaptive explicit time-stepping procedure. The time-stepping procedure is based on calculating the residual Res based on (S43). In each time-step, we calculate a possible maximum time step d*τ* by setting it inversely proportional to Res

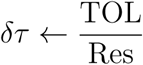

where the tolerance TOL is set beforehand. We also save the previous time step Δ*τ*

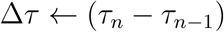

and then the new time step is calculated by using the arithmetic mean according to the following rule.

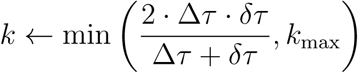

In similarity with the tolerance TOL, another numerical parameter corresponding to the maximum step length *k*_max_ is defined.

To test the validity of our implementation, a test-problem was constructed. Here we solved the homogeneous ODE-problem

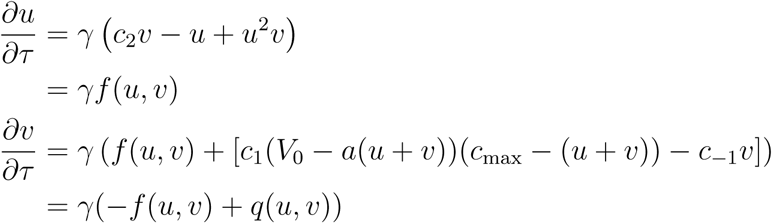

with trivial initial conditions.

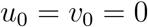

Then, compared the spatial averages of the PDE-solutions to Eq S43 given by

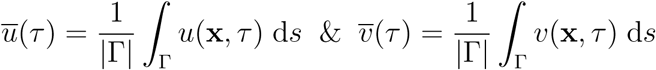

with the ODE-solutions. Again the initial conditions of *u* and *v* in the PDE-setting where set to 0 everywhere in the domain while the initial conditions of *V* was set to *V* (**x**, 0) = *V*_0_ ∀ **x** ∈ Ω. The spatial averages 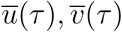 corresponded to the ODE-solutions *u*(*τ*), *v*(*τ*) for the mixed implementation (Eq (S44)) of the reaction term (Fig S3) which validates our implementation.

However, the non-linear reaction terms *f* and *q* are evaluated at the previous time node and thereby the algorithm is explicit with respect to the “complicated terms”. A consequence of this is that a large value of the relative diffusion *d* will render the solution algorithm relatively more implicit (and thus more stable) while a large value of the reaction strength *γ* will render the algorithm relatively more explicit (and thereby less stable). An advantage of the above implementation is that is faster to solve compared to a purely implicit algorithm, however a small time-step *k*_*n*_ = *τ*_*n*_ − *τ*_*n*−1_ is required to ensure stability. Moreover, the initial condition for the above time stepping procedure is set to

**Figure S3:**
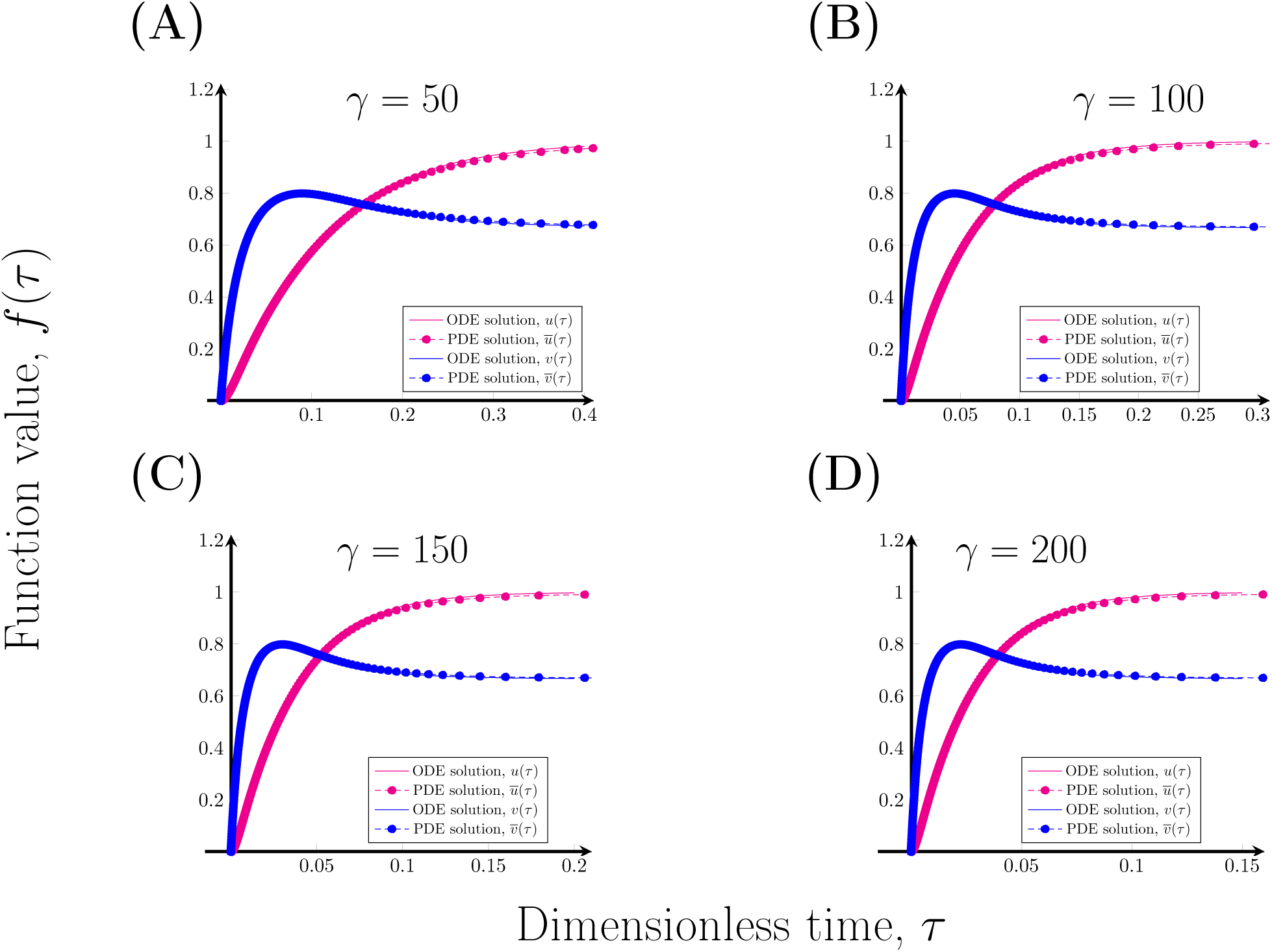
Validation of FEM-FD implementation. The states are plotted over time where the active state corresponds to the blue graph and the magenta graph to the inactive state. The ODE solutions *u, v* are represented withe whole lines while the spatial averages of the PDE solutions 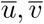 are represented by the dashed lines. The comparison is made in four cases corresponding to **(A)** *γ* = 50, **(B)** *γ* = 100, **(C)** *γ* = 150, and **(D)** *γ* = 200. The other parameters used in the simulations are: *c*_1_ = 0.05, *c*_−1_ = 0.10, *c*_1_ = 0.50, *D* = 1000, *V*_0_ = 6.0 and *c*_max_ = 3.0.

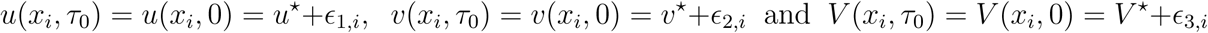

for all spatial nodes *x*_*i*_, *i* = 1, *…, N* where *N* ∈ ℕ_+_ is the total number of nodes. Above, we set the initial conditions for all spatial nodes to the steady-state value (*u*^∗^, *v*^∗^, *V* ^∗^) at which diffusion-driven instability occurs plus a noise term *ϵ*_*k,i*_ (0, *σ*), *k ϵ*{1, 2, 3 }which is normally distributed with zero mean and standard deviation *σ*. For the implementation, we have implemented a noise term of *σ* = 0.1.

The presented simulations corresponds to the numerical solutions to the problem (S40) given by this combined approach. In summary, our methodology entails applying the FEM on the above variational formulation (S43) repeatedly while advancing in time using FDs corresponding to an implicit Euler scheme. Thus, given a spatial disretisation in terms of a mesh (Fig S2B), a partitioning of the time-line *T* ([0, *τ*_max_]) = *τ*_0_, *τ*_1_, *τ*_2_, *…, τ*_*n*_, *…, τ*_max_ and initial conditions *u*_0_, *b*_0_, *ቨ*_0_ = *u* (*τ*_0_), *v* (*τ*_0_), *ቨ* (*τ*_0_) *ℌ* corresponding to the initial concentration profiles of the three states it is possible to solve the problem numerically using the finite element method based on (S43) in order to obtain the finite element solutions in each time node in *T* ([0, *τ*_max_]). Regarding the initial conditions {*u*_0_, *b*_0_, *ቨ*_0_} ∈ *ℌ*, we have initiated the three states inhomogeneously, as described above, in their respective domains (i.e. Ω for *ቨ* and Γ for *u* and *v*) with a small perturbation around each steady state value.

#### S2.3 Empirical pole-recognition algorithm

Two pole properties that we are especially interested in are pole size and time to polarisation. These properties are obtained by studying the concentration profile of active Cdc42. It is often quite easy to visually verify that a concentration profile is polarised, but to quantitatively determine when a pole arises and its extent is not trivial. We have defined mathematical condition for pole, such that a pole is reached when the concentration profile of the active Cdc42 is approximately constant, i.e. *∂u/∂t* ≈ 0, and that the maximum concentration differs significantly from the minimum.Numerically, this entails approximating the time derivative with a finite difference and then set two numerical tolerances for when these conditions are deemed to be satisfied. In an attempt to make pole-recognition more consistent, we have developed and implemented a pole-recognition algorithm. The algorithm uses tolerance parameters that have been empirically calibrated in order to reach some agreement between visual and numerical polarisation. The consistency in pole-recognition thus heavily relies on using fixed tolerance parameter values after an initial calibration phase. The algorithm checks the new discrete finite element concentration of active Cdc42, *u* (*τ*_*n*_) for every discrete time *τ*_*n*_ in the time-stepping. The main steps of the algorithm are outlined in pseudo-code below where we use *U*^*n*^ to denote the array of all *surface* nodal values of *u* (*τ*_*n*_). We let 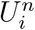 denote an arbitrary element of *U*^*n*^. The empirical tolerance parameters are all denoted TOL with some index.

##### Algorithm 1 Empirical pole-recognition (performed at discrete time *τ*_*n*_)

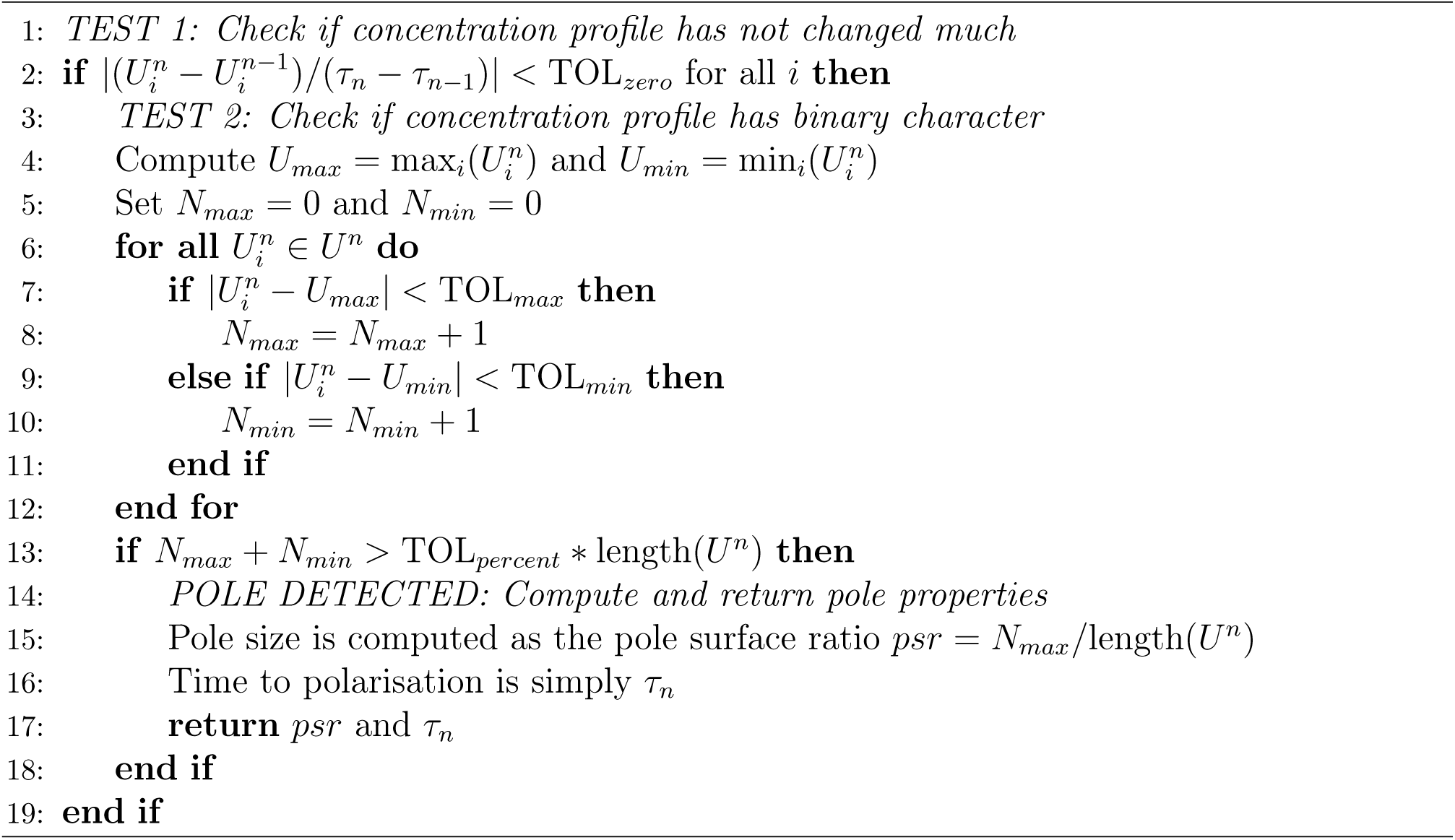

### S3 Additional results

In this section, we present additional results from the simulations of cell polarisation. Specifically, the subsequent plots focus on the effect of changing the kinetic rate parameters on the polarisation process. Also, we display the results of increasing the relative diffusion with a relative scale, that is where *u*_max_ varies for the different values of *d*.

#### S3.1 Varying the kinetic parameters

Similarly to the effect of changing the relative diffusion *d* (Fig 4 on page 12) and the relative reaction strength *γ* (Fig 6 on page 13), the effect of changing the kinetic parameters on the cell polarisation process can be investigated. Therefore, the results of the simulations for different kinetic parameters in the (*c*_−1_, *c*_2_)-plane (Fig S4) and the (*c*_1_, *c*_2_)-plane (Fig S5) have been visualised.

**Figure S4:**
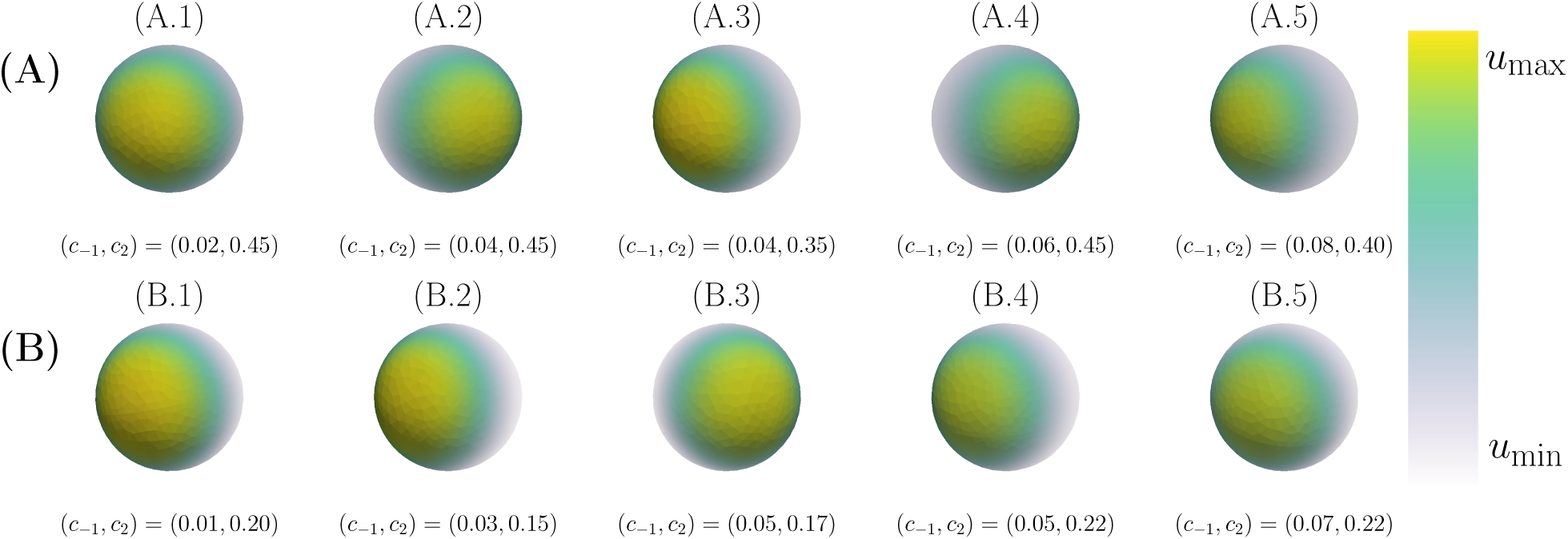
Final patterns for different kinetic parameters within the (c_−1_, c_2_)-plane. The final patterns for various parameters in the (*c*_−1_, *c*_2_)-plane are displayed in two cases, namely classic and non-classic diffusion driven instability. In both cases, the final time when the pattern is formed *τ*_final_ and the maximum and minimum concentration of active Cdc42 *u*_max_ and *u*_min_ are calculated as functions of the kinetic rate parameters. **(A)** *Classic*: (A.1): (*c*_−1_, *c*_2_, *τ*_final_, *u*_max_, *u*_min_) = (0.02, 0.45, 4.40, 3.77, 0.38), (A.2): (*c*_−1_, *c*_2_, *τ*_final_, *u*_max_, *u*_min_) = (0.04, 0.45, 4.75, 3.64, 0.40), (A.3): (*c*_−1_, *c*_2_, *τ*_final_, *u*_max_, *u*_min_) = (0.04, 0.35, 3.61, 3.83, 0.29), (A.4): (*c*_−1_, *c*_2_, *τ*_final_, *u*_max_, *u*_min_) = (0.06, 0.45, 5.42, 3.55, 0.41) and (A.5) (*c*_−1_, *c*_2_, *τ*_final_, *u*_max_, *u*_min_) = (0.08, 0.40, 4.95, 3.57, 0.37). **(B)** *Non-classic*: (B.1): (*c*_−1_, *c*_2_, *τ*_final_, *u*_max_, *u*_min_) = (0.01, 0.20, 5.50, 4.26, 0.15), (B.2): (*c*_−1_, *c*_2_, *τ*_final_, *u*_max_, *u*_min_) = (0.03, 0.15, 3.38, 4.18, 0.11), (B.3): (*c*_−1_, *c*_2_, *τ*_final_, *u*_max_, *u*_min_) = (0.05, 0.17, 6.61, 4.06, 0.13), (B.4): (*c*_−1_, *c*_2_, *τ*_final_, *u*_max_, *u*_min_) = (0.05, 0.22, 4.70, 3.92, 0.18) and (B.5) (*c*_−1_, *c*_2_, *τ*_final_, *u*_max_, *u*_min_) = (0.07, 0.22, 4.05, 3.91, 0.18). In both cases, the overall parameters are: *c*_1_ = 0.05, *V*_0_ = 6.0, *c*_max_ = 3.0, *a* = 3, *d* = 10 and *γ* = 25.

**Figure S5:**
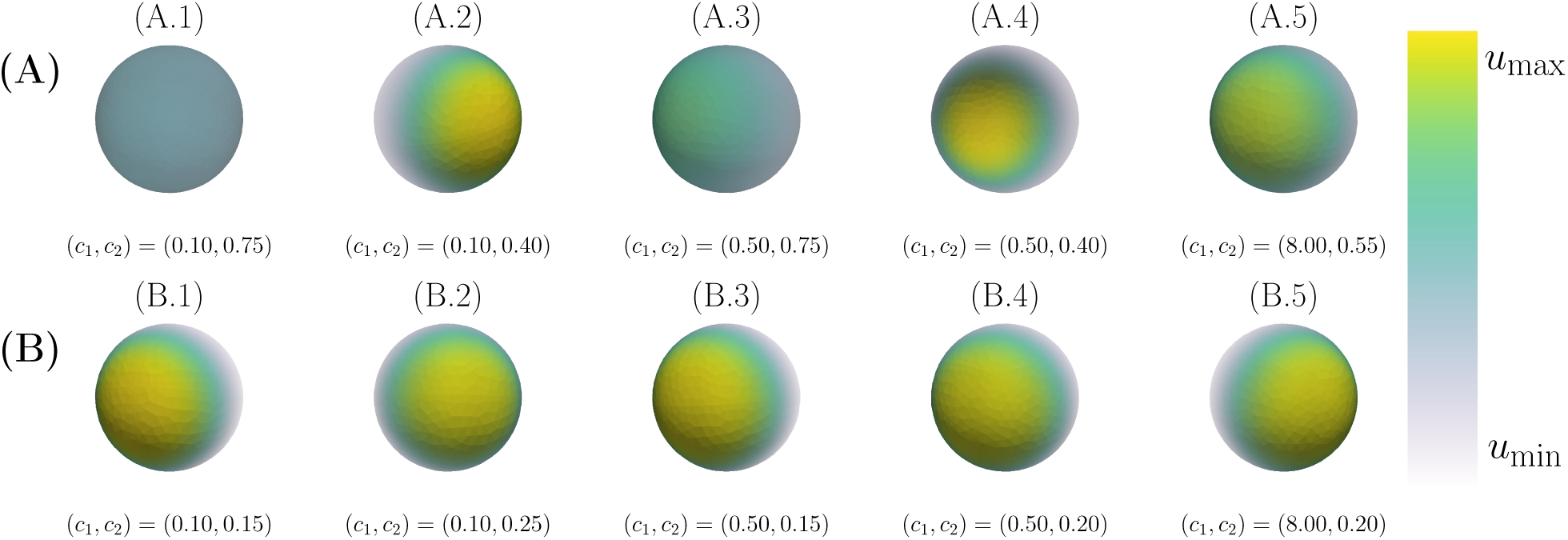
Final patterns for different kinetic parameters within the (c_1_, c_2_)-plane. The final patterns for various parameters in the (*c*_1_, *c*_2_)-plane are displayed in two cases, namely classic and non-classic diffusion driven instability. In both cases, the final time when the pattern is formed *τ*_final_ and the maximum and minimum concentration of active Cdc42 *u*_max_ and *u*_min_ are calculated as functions of the kinetic rate parameters. **(A)** *Classic*: (A.1): (*c*_1_, *c*_2_, *τ*_final_, *u*_max_, *u*_min_) = (0.10, 0.75, 10.59, 1.54, 1.28), (A.2): (*c*_1_, *c*_2_, *τ*_final_, *u*_max_, *u*_min_) = (0.10, 0.40, 6.69, 3.65, 0.35), (A.3): (*c*_1_, *c*_2_, *τ*_final_, *u*_max_, *u*_min_) = (0.50, 0.75, 19.0, 2.65, 0.89), (A.4): (*c*_1_, *c*_2_, *τ*_final_, *u*_max_, *u*_min_) = (0.50, 0.40, 6.48, 3.59, 0.36) and (A.5) (*c*_1_, *c*_2_, *τ*_final_, *u*_max_, *u*_min_) = (8, 0.55, 6.51, 3.27, 0.54). **(B)** *Non-classic*: (B.1): (*c*_1_, *c*_2_, *τ*_final_, *u*_max_, *u*_min_) = (0.10, 0.15, 5.08, 4.02, 0.12), (B.2): (*c*_1_, *c*_2_, *τ*_final_, *u*_max_, *u*_min_) = (0.10, 0.25, 2.88, 3.88, 0.20), (B.3): (*c*_1_, *c*_2_, *τ*_final_, *u*_max_, *u*_min_) = (0.50, 0.15, 8.38, 3.93, 0.12), (B.4): (*c*_1_, *c*_2_, *τ*_final_, *u*_max_, *u*_min_) = (0.50, 0.20, 7.97, 3.87, 0.16) and (B.5) (*c*_1_, *c*_2_, *τ*_final_, *u*_max_, *u*_min_) = (8, 0.20, 9.04, 3.86, 0.16). In both cases, the overall parameters are: *c*_−1_ = 0.05, *V*_0_ = 6.0, *c*_max_ = 3.0, *a* = 3, *d* = 10 and *γ* = 25.

#### S3.2 Effect of increasing diffusion with an absolute scale

Visually, It is not clear that the maximum concentration of active Cdc42 *u*_max_ increases with an increasing relative diffusion *d* in Fig 4 on page 12 of the article. In this plot, the concentration scale, i.e. the colourbar, is relative where the value of *u*_max_ varies between the different cases. The advantage with this representation is that it is visually clear that the relative size of the pole decreases with an increasing relative diffusion. Although, the decrease in *u*_max_ is presented in the figure caption it is also possible to visually present this effect. To this end, the same figure has been reproduced with an absolute scale on the colour bar (Fig S6).

**Figure S6:**
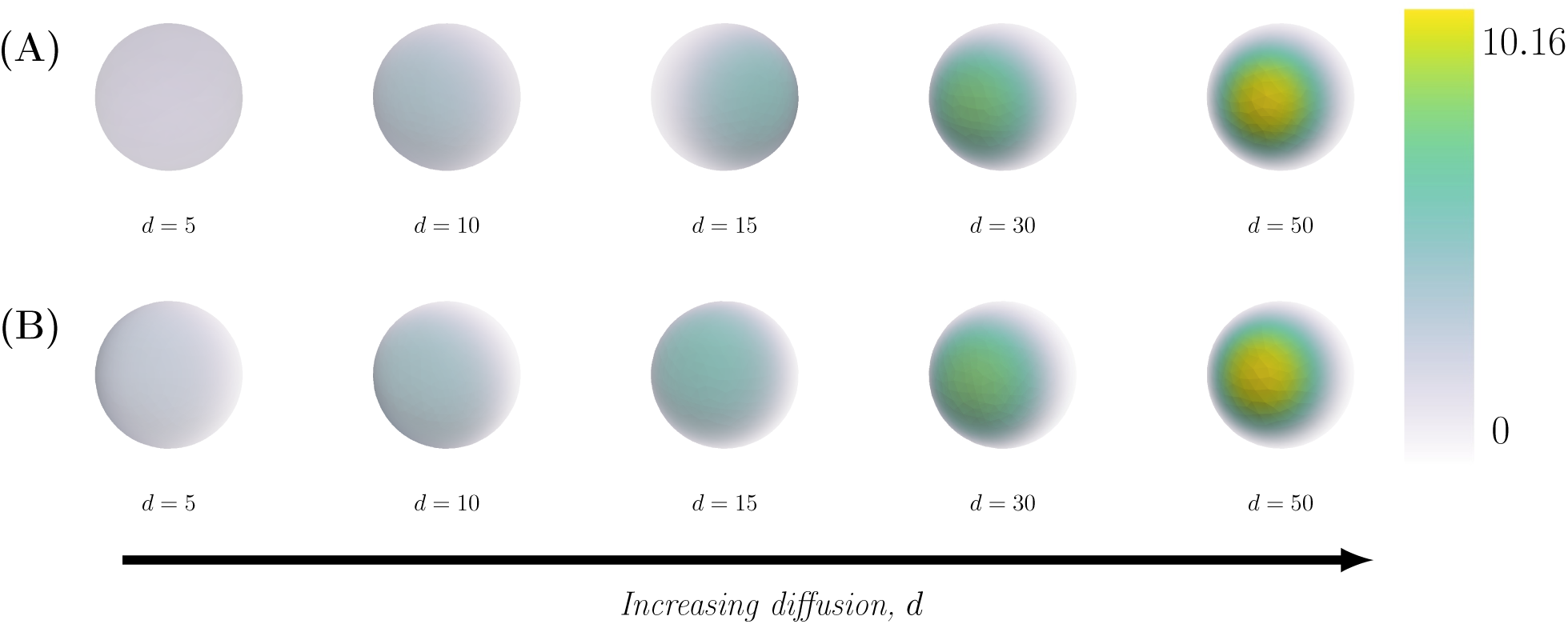
Final patterns for increasing relative diffusion with absolute scale. The final patterns for increasing relative diffusion *d* are displayed in two cases, namely classic and non-classic diffusion driven instability. In both cases, the final time when the pattern is formed *τ*_final_ and the maximum and minimum concentrations of active Cdc42 *u*_max_ and *u*_min_ are calculated as functions of the kinetic rate parameters. **(A)** *Classic*: The overall parameters are (*c*_1_, *c*_−1_, *c*_2_) = (0.05, 0.04, 0.45) with specific parameters (from left to right): no pattern is formed for (*d, τ*_final_, *u*_max_, *u*_min_) = (5, 15, 20.89, 1.34, 1.11), (*d, τ*_final_, *u*_max_, *u*_min_) = (10, 4.44, 3.65, 0.40), (*d, τ*_final_, *u*_max_, *u*_min_) = (15, 3.65, 4.85, 0.30), (*d, τ*_final_, *u*_max_, *u*_min_) = (30, 2.65, 7.42, 0.20) and (*d, τ*_final_, *u*_max_, *u*_min_) = (50, 2.17, 9.83, 0.15). **(B)** *Non-classic*: The overall parameters are (*c*_1_, *c*_−1_, *c*_2_) = (0.05, 0.03, 0.15) with specific parameters (from left to right): (*d, τ*_final_, *u*_max_, *u*_min_) = (5, 4.0, 2.77, 0.17), (*d, τ*_final_, *u*_max_, *u*_min_) = (10, 4.38, 4.18, 0.11), (*d, τ*_final_, *u*_max_, *u*_min_) = (15, 2.87, 5.29, 0.09), (*d, τ*_final_, *u*_max_, *u*_min_) = (30, 1.95, 7.77, 0.06) and (*d, τ*_final_, *u*_max_, *u*_min_) = (50, 1.96, 10.166, 0.04). In both cases, the overall parameters are: *V*_0_ = 6.0, *c*_max_ = 3.0, *a* = 3 and *γ* = 25.

## Footnotes

S1 The spherical case is analogous.

S2 A closely connected concept is that of fundamental solutions which are also expressed in terms of these polynomials. For a specific example of what the fundamental solutions to the heat equation on the unit sphere look like in the case of rotational invariance, see [S43].

S3 The critical point *u*_0_ is a local max-point of the function *v*(*u*).

S4 The (*c*_−1_, *c*_2_)-space is generated completely analogously.

